# *Photorhabdus* Virulence Cassettes: extracellular multi-protein needle complexes for delivery of small protein effectors into host cells

**DOI:** 10.1101/549964

**Authors:** Isabella Vlisidou, Alexia Hapeshi, Joseph R. J. Healey, Katie Smart, Guowei Yang, Nicholas R. Waterfield

## Abstract

*Photorhabdus* is a highly effective insect pathogen and symbiont of insecticidal nematodes. To exert its potent insecticidal effects, it elaborates a myriad of toxins and small molecule effectors. Among these, the *Photorhabdus* Virulence Cassettes (PVCs) represent an elegant self-contained delivery mechanism for diverse protein toxins. Importantly, these self-contained nanosyringes overcome host cell membrane barriers, and act independently, at a distance from the bacteria itself. In this study, we demonstrate that Pnf, a PVC needle complex associated toxin, is a Rho-GTPase, which acts via deamidation and transglutamination to disrupt the cytoskeleton. TEM and Western blots have shown a physical association between Pnf and its cognate PVC delivery mechanism. We demonstrate that for Pnf to exert its effect, translocation across the cell membrane is absolutely essential.

**SIGNIFICANCE STATEMENT:** Here we provide an up to date analysis of the nano-scale syringe-like molecular devices that *Photorhabdus* use to manipulate invertebrate hosts, the PVC system. They are related to the *Serratia* Anti-Feeding Prophage and the Psuedoalteromonas MAC system. All these systems are in turn more distantly related to the well characterized Type VI secretion system currently receiving a great deal of attention. We demonstrate for the first time that the PVC nanosyringes are physically “loaded” with an effector protein payload before being freely released. The PVCs therefore represent bacterial molecular machines that are used as “long-range” protein delivery systems. This widespread class of toxin delivery system will likely prove of great significance in understanding many diverse bacteria/host interactions in future.

## INTRODUCTION

Bacteria belonging to the Enterobacteriacae genus *Photorhabdus* exist in a symbiotic partnership with entomopathogenic *Heterorhabditis* sp. nematodes. This Entomopathogenic Nematode complex (EPN) comprises a highly efficient symbiosis of pathogens that is commonly used as a biological agent to control crop pests [1]. The *Photorhabdus* bacteria are delivered into the hemocoel of the insect, after regurgitation from the worm, where they resist the insect immune response and rapidly kill the host via septicaemic infection. Insect tissues are subsequently bio-converted into a dense soup of *Photorhabdus* bacteria, which provide a food source to support the replication of the nematode. As food resources are depleted *Photorhabdus* re-associates with infective juvenile nematodes, and together they emerge from the insect cadaver able to re-infect a new host [2, 3]. Three major species have been formally recognized to date within the genus -*P. luminescens, P. asymbiotica*, and *P. temperata*. It should be noted however that with increasing numbers of *Photorhabdus* genome sequences becoming available, the genus structure is under revision [4]. In addition to the normal insect life cycle, *P*. *asymbiotica* is also the etiological agent of a serious human infection termed *Photorhabdosis*, which is associated with severe ulcerated skin lesions both at the initial infection foci and later at disseminated distal sites [5-8].

The *Photorhabdus* genome encodes a diverse repertoire of virulence genes encoding for protein toxins, proteases and lipases for combating diverse hosts, that can be found in chromosomally encoded pathogenicity islands [9-14]. In addition the bacteria also secrete a potent cocktail of other biologically active small molecules to preserve the insect cadaver in the soil from competing saprophytes and microbial predators such as amoeba [15, 16]. Several classes of *Photorhabdus* protein insecticidal toxins have now been well characterised including the Toxin Complexes [17-24], the binary PirAB toxins [25-27] and the large single polypeptide Mcf (“makes caterpillars floppy”) toxins [28-30].

A fourth class of highly distinct toxin delivery systems first identified in *Photorhabdus* are the “*Photorhabdus* virulence cassettes”, or PVCs [31]. These represent operons of around 16, conserved, structural and synthetic genes (from hereon just described as the structural genes) encoding for a phage “tailocin” like structure [32] and one or more tightly linked downstream toxin-effector like genes. Genomic analysis of multiple strains of *Photorhabdus* revealed they often encode up to five or six copies of the operon, each with unique downstream effector genes [33].

It should be noted that PVC-like elements are not restricted to *Photorhabdus* as a well-characterized homologous operon can also be found on the pADAP plasmid of the insect pathogenic bacteria *Serratia entomophila* [34]. This system has been named the anti-feeding prophage (AFP), as it is responsible for the cessation of feeding in the New Zealand grass grub host. Recent cryo-electron microscopy studies have revealed that, morphologically, AFP resembles a simplified version of the sheathed tail of bacteriophages such as T4, including a baseplate complex. It also shares features with type-VI secretion systems, with the central tube of the structure having a similar diameter and axial width to the Hcp1 hexamer of *P. aeruginosa* T6SS [35]. One important difference between the PVC and T6SS machinery is that the T6SS relies upon direct contact between host and bacterial cell, and is anchored in to the membrane by a substantial membrane complex whose structure is still being elucidated [36], whereas the PVC needle complex is freely released into the surrounding milieu and so can act at a distance.

Furthermore, recent reports have indicated that other more diverse bacteria can also make similar needle complexes for manipulation of eukaryotic hosts. A well-characterized example is the production of analogous devices by the marine bacterium *Pseudoalteromonas luteoviolacea* (Figure 1A). These structures are involved in the developmental metamorphosis of the larvae of the tubeworm *Hydroides elegans*, and they are deployed in outward-facing arrays comprising about 100 contractile structures, with baseplates linked by tail fibres in a hexagonal net [37]. Interrogation of sequence databases with PVC protein sequences suggests many other more diverse tailocin-like systems are yet to be characterized [38]. These include operons closely related to the PVCs in *Xenorhabdus bovienii* CS03, *Yersinia ruckeri* ATCC29473 and *Vibrio campbellii* AND4. In addition, evidence of more diverse elements, like that of *P. luteoviolacea*, can also be seen. To address this, we have recently performed an exhaustive analysis of all available prokaryotic and archaeal genome sequences in the public databases to look at the distribution of *pvc*-like elements (unpublished data). This suggests that PVC-like nano-syringes and their distant cousins are of enormous ecological and perhaps biomedical significance.

**Figure 1.**
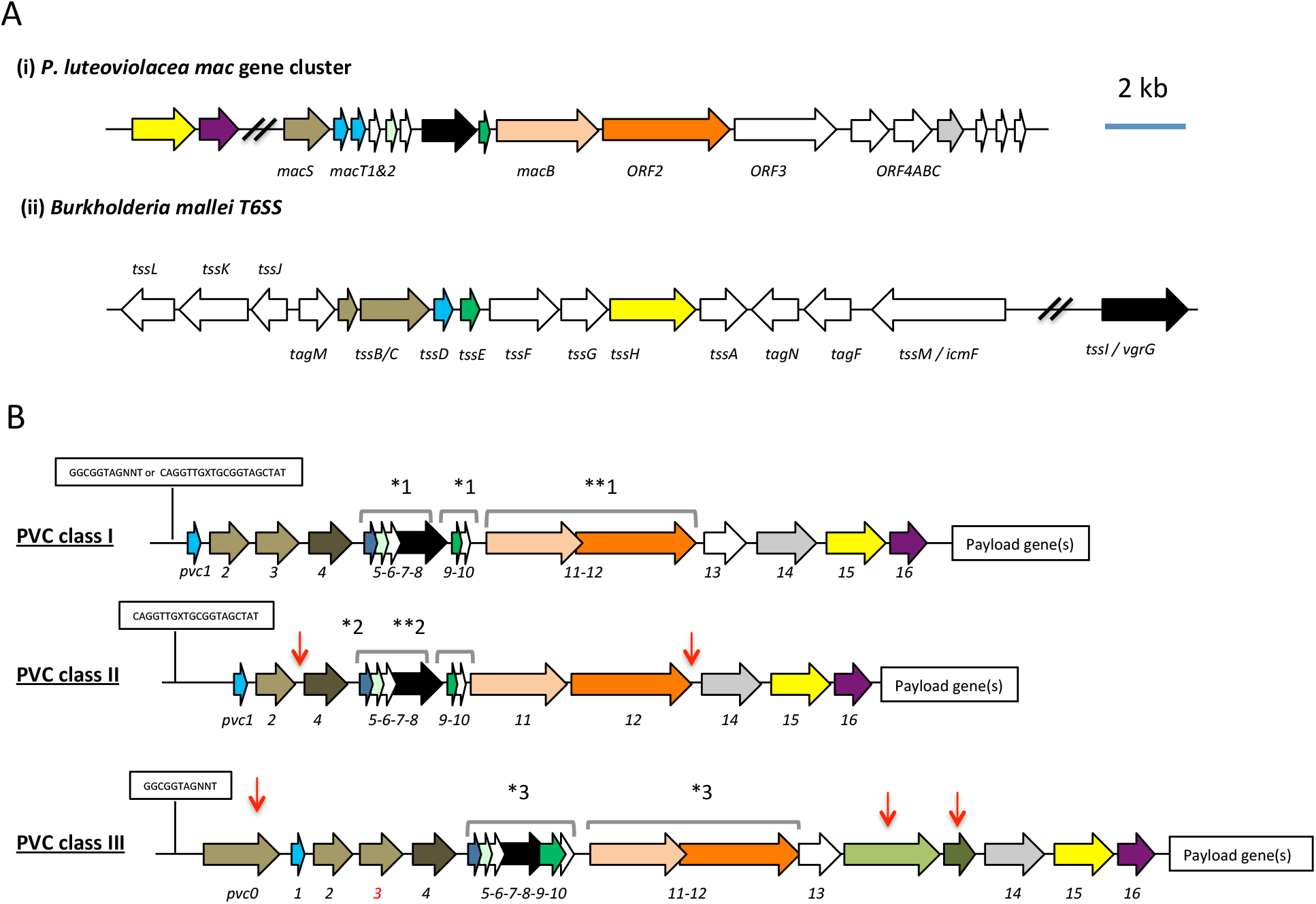
(**A**) Similarity between PVCs and two diverse protein secretion systems, (i) the *P. luteoviolacea mac* gene cluster and (ii) The type-VI secretion system (T6SS) from *Burkholderia mallei.* Homologous protein sequences are coloured coded. **(B)** Three classes of PVC structural operons observed in the genomes of *Photorhabdus* and members of other genera. Types 1-3 are exemplified by <u>PVC</u>*<u>pnf</u>*<u>, PVC</u>*<u>lopT</u>* <u>and PVC</u>*<u>PaTox</u>* respectively. Homologous genes are colour coded. Red arrows represent variations relative to the representative type I PVC*pnf* operon of *P. asymbiotica* ATCC43949, *pvc1* (PAU_03353) to *pvc16* (PAU_03338). Predicted functions of individual Pvc proteins based on homology to known proteins can be seen in FigS1C. The boxed “GGCGGTAGNNT” or “CAGGTTGXTGCGGTAGCTAT” sequences represent positions of the conserved RfaH anti-termination protein and cryptic operator sequences respectively. Square brackets above certain genes indicate apparent translational coupling. More specifically; *1 indicates coupling in PVC*pnf* and PVC*cif* of *Pl*^TT01^, *Pa*^ATCC43949^, *Pa*^PB68^ and *Pa*^Kingscliff^ and in the *Serratia entomophila afp* operon in addition to an uncharacterised PVC in *Yersinia ruckeri* ATCC 29473. **1 indicates these genes are not coupled in *Pa*^Ki^^ngscliff^. *2 indicates coupling in PVC*lopT* of *Pl*^TT01^, *Pa*^ATCC43949^, *Pa*^PB68^ and *Pa*^Kingscliff^. **2 indicates these genes are not coupled in *Pa*^Kingscliff^. *3 indicates coupling in PVC*Patox* of *Pa*^ATCC43949^and *Pa*^Kingscliff^ (although *pvc11* possibly contains a frame-shift in *Pa*^Kingscliff^). The *pvc3* is also deleted in *Pa*^Kingscliff^.

Here we focus on a single *Photorhabdus pvc* operon (which elaborates the PVC*pnf* needle complex [31] to understand the relationship between the structural genes and the tightly linked effector gene, *pnf*. We confirm *in vivo* expression during insect infection and reveal a high level of population heterogeneity of expression *in vitro*. We demonstrate for the first time the physical association of the Pnf effector toxin protein with the secreted structural needle complex using Western blot and electron microscopy. Furthermore, we prove that the cognate Pnf effector needs to be delivered into the eukaryote cell cytoplasm to exert any measurable effect and confirm its predicted activity targeting small Rho-GTPase target proteins. Taken together this work describes an important new class of protein toxin secretion and injection delivery systems which, unlike the well-described Types III, IV and VI systems, can act “at a distance”, requiring no intimate contact between bacteria and host cells.

## RESULTS

### A bioinformatic analysis of *pvc* structural operon sequences

A comparison of *pvc* structural operons identified in the genome sequences of *Photorhabdus* and certain members of other genera, available at the time of publication ([12, 14, 39] and our unpublished data), allowed us to define three distinct genetic sub-types. The PVC*pnf* operon belongs to class I, which has 16 structural genes and three translationally coupled gene blocks, and is of the type typically seen in non-*Photorhabdus* genera. Class II and III operons differ in the number of structural genes and translationally coupled gene blocks (Figure 1B). Given the diversity of *pvc*-operons, and their typically poor annotation in genome sequences, it is necessary here to define a nomenclature protocol to allow reference to any given operon. An example of the method we have adopted is as follows; [*Pa* ^ATCC43949^ PVC*pnf*], where *Pa* ^ATCC43949^ is species and strain, in this case *Photorhabdus asymbiotica* strain ATCC43949 and PVC*pnf* is the specific operon within that genome with the suffix referring to one of the tightly linked effectors, in this case the *pnf* effector gene. We will also include gene identifiers for either end of the operon where appropriate, which in this case would be PAU_03353-PAU_03332, which are the genes for *pvc1* and *pnf* respectively.

With reference to published literature and a detailed bioinformatic analysis of promoter regions upstream of the *pvc1* genes, we can identify two distinct, potential *cis*-operator sequences. Firstly operons belonging to classes I (e.g. PVC*pnf*) and III (e.g. PVC*PaTox*) typically encode the highly conserved RfaH operator sequence, GGCGGTAGNNT [40]. It is possible that more degenerate RfaH operator sequences exist in other operons although this remains unclear. Secondly, all class II operons (e.g. PVC*lopT*) and certain class I operons (e.g. PVC*units*1-4) encode a minimal cryptic conserved sequence motif, CAGGTTGXTGCGGTAGCTAT. In both cases these conserved *cis*-encoded sequences are located between the *pvc1* gene and the transcription start sites, as defined by previous RNA-seq analysis ([41] and unpublished data).

Several observations suggest that horizontal gene transfer has been responsible for the dissemination of many observed *pvc*-operons. These include; the location of the *S. entomophila afp* on a horizontally transmissible plasmid, the presence of four *pvc* operons in tandem in *P. luminescens* TT01 (directly adjacent to a type IV DNA conjugation pilus operon), the presence of multiple *pvc* operons in any given genome and the suggestion that several operons are regulated by RfaH. While there is no experimental evidence to confirm an exact mechanism by which this may occur, a closer inspection of the sequences flanking [*Pl*^TT01^PVC*u4*] suggests that at least this operon was acquired as a composite transposon. Remnants of insertion sequence (IS) elements can be seen flanking this operon, with only the outer inverted repeats remaining intact [TTATATTGAA(t/g)GAATATTAAGCAAGAAAC], and YhgA-like IS transposase genes belonging to the (transposase_31 superfamily) still associated with both the 5’ and 3’ flanks of the PVC*u4* operon. It should be noted that IS element remnants could also be seen flanking many other *pvc* operons suggesting that IS dependant transposition has been a common mechanism involved in *pvc* horizontal dissemination. However, our own phylogenetic studies suggest that *pvc*-operons have been co-evolving with their host genomes for some time, indicating that horizontal transfer is likely the method of original acquisition, but may not be as active presently. This is supported by the fact that an automatic prediction of horizontal gene transfer regions (HGTs) using Alien Hunter 1.7 [42] either did not detect any HGT elements spanning the structural regions of PVCs or in the cases where such an element was detected it was assigned a low confidence score (Figure S2).

An analysis of the conservation of individual genes across different *pvc* operons at both DNA and protein sequence levels suggests that either recombination or diversifying selection is more likely to have occurred in the more 3’ regions of the operons (Figure S1A). This is perhaps no surprise as each *pvc* operon can be seen to encode different effector genes in the 3’ payload region of the operons. An analysis of conservation of protein sequences of the *pvc* operons showed that within *pvc*-operons a good deal of variability is possible while presumably retaining the ability to produce a similar macromolecular structure (Figure S1B). This is supported by HHPRED structural homology comparisons for equivalent PVC proteins across different operons, despite often-variable primary amino acid sequences (data not shown). We note that the most diverse protein seen in *pvc*-operons is that of the predicted tail fibre proteins, Pvc13, which we may expect if different *pvc*-operons are adapted for different host cell targets. Paralogous genes within *pvc*-operons include *pvc1* and *pvc5* which encode homologs of Hcp, the inner tube protein of contractile tube mechanisms such as T6SS and phage protein Gp27 and *pvc2*, -*3* and -*4* which encode homologues of the outer sheath proteins of phage [43] and T6SS [44]. Figure S1C illustrates the organisation of the [*Pa*^ATCC43949^ PVC*pnf*] operon used as a model system in our experimental studies described here, showing the top HHPRED structural homology hits and predicted roles for each encoded protein at the time of writing.

### A bioinformatic analysis of *pvc*-operon effector gene sequences

A comparison of the 3’ effector “payload regions” of different *pvc* operons reveals a large diversity of effector genes, with a range of predicted activities, covering a large range of sizes and isoelectric point values (data not shown). Some operons encode only a single putative effector, e.g. [*Pa*^ATCC43949^ PVC*PaTox* PAU_02249-02230] while others have several, either tandem homologues of one another, e.g. [*Pa*^ATCC43949^ PVC*u4* PAU_02790-02808] or entirely unrelated putative effector genes, e.g. [*Pa*^ATCC43949^ PVC*lopT* PAU_02112-02095]. Many effector genes are also tightly linked to transposase gene remnants suggesting they are typically exchanged by horizontal acquisition. This is further supported by the observation that orthologous *pvc*-operons in the same chromosomal context may have different effector genes in different strains. A good example of this being the unrelated effector genes seen in the orthologous structural “PVC*pnf*” operon loci of *Pa*^*Kingscliff*^ and *Pa*^ATCC43949^ which carry a tyrosine glycosylase and Pnf (this paper) respectively. Analysis with Alien Hunter 1.7, suggests that certain *pvc*-operon / effector associations are ancestral to any given species. For example the association of the *pvc17* effector with PVC*u4*, and the multiple linked effectors with the PVC*lopT* operon in both *Pa*^ATCC43949^ and *Pl*^TT01^. Conversely other *pvc*-operons show evidence of recent horizontal acquisition of their 3’linked effectors, e.g. PVC*cif* and PVC*pnf* (not shown)

### Expression of PVC *pnf in vitro* and *in vivo*

A previous RNA-seq analysis of global transcription in three strains; *P. asymbiotica* ^ATCC43949^ [41], *P. asymbiotica* ^Kingscliff^ and *P. luminescens* ^TT01^ (unpublished) showed condition dependent expression of certain *pvc*-operons but not all. Therefore, due to the diversity of *pvc* operons and effectors in *Photorhabdus*, we focused on a single model class I *pvc* operon, [*Pa*^ATCC43949^ PVC*pnf*], to elucidate the relationship between the conserved structural and effector proteins. This operon was selected as it elaborates a well-defined needle complex structure (as observed by electron microscopy) which has potent insect killing activity when heterologously expressed in *E. coli* [31]. This operon has two putative effector genes in the downstream “payload region”, PAU_03337, which shows similarity to adenylate cyclase toxins (e.g. the anthrax Edema Factor and Pseudomonas ExoY toxin) and *pnf* (PAU_03332). While the predicted activity of PAU_03337 has not been tested directly, when expressed in the NIH-3T3 cell cytoplasm (in transient transfection experiments) it did produce a highly unusual cytoskeleton phenotype [31]. Pnf (<u>P</u>hotorhabdus <u>n</u>ecrosis factor) is a homologue of the active site domain of the *Yersinia* CNF2 (Cyto Necrosis Factor 2) toxin, which has small-GTPase deamidase and transglutaminase activities [45].

In order to confirm the expression of this model *pvc*-operon in *Photorhabdus* during an insect infection we constructed transcription-translation reporter plasmids in which the promoter regions and the first 150 bp of coding sequence of *pvc1, pnf* [both from *Pa*^ATCC43949^ PVC*pnf*] and the *P. asymbiotica* chromosomal *rpsM* ribosomal “housekeeping” gene (as a positive control) were genetically fused in frame to a *gfpmut*2 gene with no start codon (referred to hereon as *pvc1*::*gfp, pnf*::*gfp* and *rpsM*::*gfp* reporters). Note, the genomic context and our previous unpublished RT-PCR studies suggested that *pnf* had its own promoter and could be transcribed independently of the *pvc* structural genes. As we are unable to transform *Pa*^ATCC43949^ itself, these plasmids were transformed into the well-characterised and genetically tractable strain *P. luminescens*^TT01^ to provide suitable reporter strains for *in vitro* and *in vivo* expression studies. For *in vitro* studies we cultured the bacteria in LB medium supplemented with *Manduca sexta* clarified hemolymph and grown to late stationary phase, before microscopic examination. For *in vivo* studies, we injected the reporter strains into *M. sexta*, and allowed the infection to establish before macroscopic examination of insect tissues *in situ* using a (fluorescence) dissecting microscope. We also took hemolymph samples from these insects and visualised the hemocytes and bacteria microscopically using confocal microscopy.

Figure 2A shows expression of GFP reporter from the *rpsM* positive control and both the *pvc1*::*gfp* and *pnf*::*gfp* reporters in LB supplemented with *M. sexta* hemolymph, although not in all cells of the bacterial population (see below). Furthermore, we also saw expression in bacteria in the *ex vivo* hemolymph samples taken during infection of live insects (Figure 2A). It was also possible to confirm expression of *pnf*::*gfp* in bacteria attached to the insect trachea in localised putative biofilm masses. In this case, while the expected insect melanisation immune response could be seen to have occurred elsewhere on the trachea, it was notably absent from the *pnf* expressing bacterial biomass (Figure 2B). In order to corroborate the observations made using the plasmid based reporter constructs in *P. luminescens* ^TT01^ we also performed RT-PCR analysis of transcription of the PVC*pnf* chromosomal operon in the original *Pa*^ATCC43939^ strain. This confirmed transcription across the operon *in vitro* when the bacteria were grown at either 28°C or 37°C, although transcription of certain genes was difficult to detect *in vivo* during *Manduca sexta* infections (Figure S3).

**Figure 2.**
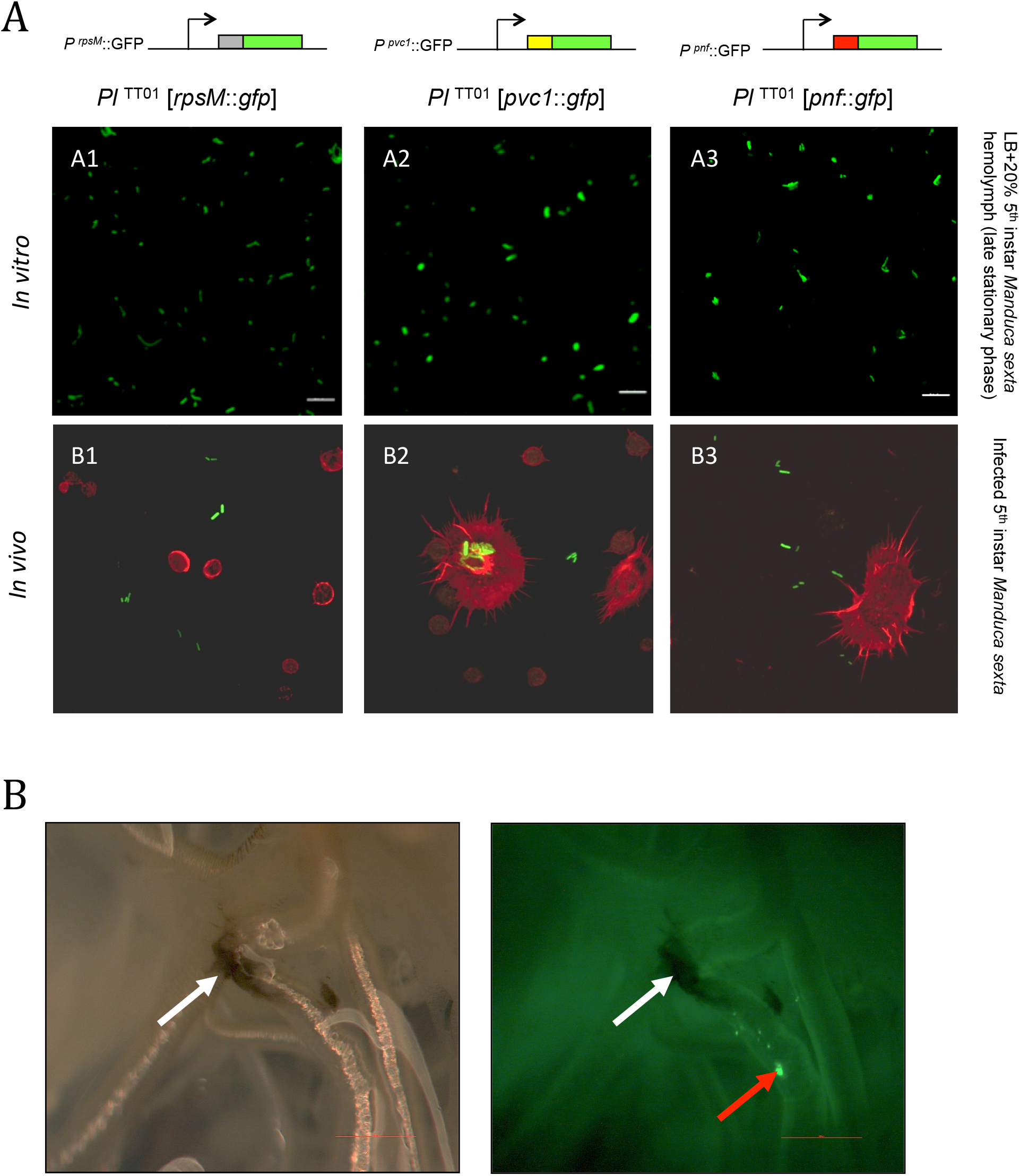
(**A**) Confocal micrographs of showing in vitro (A1-A3) and in vivo (B1-B3) expression in *Pl*^TT01^ of *gfp* transcription-translation reporter constructs of *Pa*^ATCC43949^ PVC*pnf* operon gene promoters. These plasmid-based reporters were constructed by fusing the transcription promoter regions and the first 37 codons of the target gene in frame with the second codon of *gfp*. Target gene promoters shown are (A1 and B1) the constitutively expressed *rpsM* gene, (A2 and B2) the *Pa*^ATCC43949^ PVC*pnf pvc1* structural gene and (A3 and B3) *Pa*^ATCC43949^ PVC*pnf pnf* payload toxin gene. The *in vitro* panels (A1-3) show reporter expression after growth in LB supplemented with 20% (v/v) 5^th^ instar *M. sexta* hemolymph at late stationary phase. The *in vivo* panels show *ex vivo* hemolymph from 5^th^ instar *M. sexta* infected with *Pl*^TT01^ harbouring the three different reporter constructs. The hemocyte cytoskeletons are stained red with TRITC-Phalloidin conjugate. (**B**) White light (left) and fluorescence illumination (right) of the trachea of a dissected 5^th^ instar *M. sexta* previously infected with *Pl*^TT01^ harbouring the *Pa*^ATCC43949^ PVC^pnf^ *pnf::gfp* reporter construct. Brightly fluorescent green bacteria were detected in association with the trachea (red arrow) in close proximity to melanotic nodules (white arrows), demonstrating the induction of the *pnf* promoter and the production of the Pnf::GFP fusion *in situ*. Bars show 0.1mm.

We subsequently expanded this analysis to include a transcription-translation reporter plasmid for the promoter and *pvc1* gene of an orthologue of PVC*pnf* from a different *P. asymbiotica* strain, [*Pa*^PB68^ PVC*pnf*]. In this case, we used fluorescence microscopy to assess the expression pattern across the growth phases of the originator *Pa*^PB68^ strain harbouring the reporter plasmid, when grown in LB with aeration and maintaining plasmid marker selection. Interestingly we observed a high level of population heterogeneity in expression with only very few cells expressing GFP at any one time (Figure S4). A similar level of heterogeneity in expression was also seen for reporter constructs from seven other *pvc*-operons from both *Pa*^PB68^ and *Pl*^TT01^ (data not shown). We also assessed expression in biofilms grown statically on glass slides and observed the same pattern, though with even fewer cells seen to express GFP (data not shown).

### The Pnf effector protein is physically associated with the PVC needle complex

We investigated if the Pnf effector protein actually becomes physically associated with the *pvc*-encoded needle complex we had previously visualised by electron microscopy [31]. To do this we raised anti-peptide antibodies against synthetic peptides representing amino acids 206-219 of Pnf (TGQKPGNNEWKTGR) and amino acids 130-143 (DGPETELTINGAEE) of predicted outer sheath protein Pvc2. Previously we used 2D-SDS PAGE analysis of PVC*pnf* needle complex produced by an *E. coli* cosmid clone to confirm the presence of Pvc2, along with Pvc1, 3 5, 11, 14 and 16 proteins ([31] and unpublished data). We confirmed specificity of the Pnf antibody using western blot analysis of extracts of *E. coli* heterologously expressing Pnf alone.

We first used the anti-Pnf peptide antibody to test for the presence of Pnf protein in supernatants from the native bacterial strain *Pa*^ATCC43949^. We tested for the presence of Pnf in needle complex enriched particulate preparations and clarified supernatants. We could detect Pnf in preparations enriched for the complexes but not in clarified supernatants. More specifically, the Pnf protein could only be detected in the needle complex fraction, if it was first either chemically or physically disrupted before electrophoresis (Figure 3B). Taken together these findings are consistent with the hypothesis that the Pnf protein is sequestered inside the needle complex or in some other configuration such that the TGQKPGNNEWKTGR epitope is physically hidden from access by the antibody.

**Figure 3.**
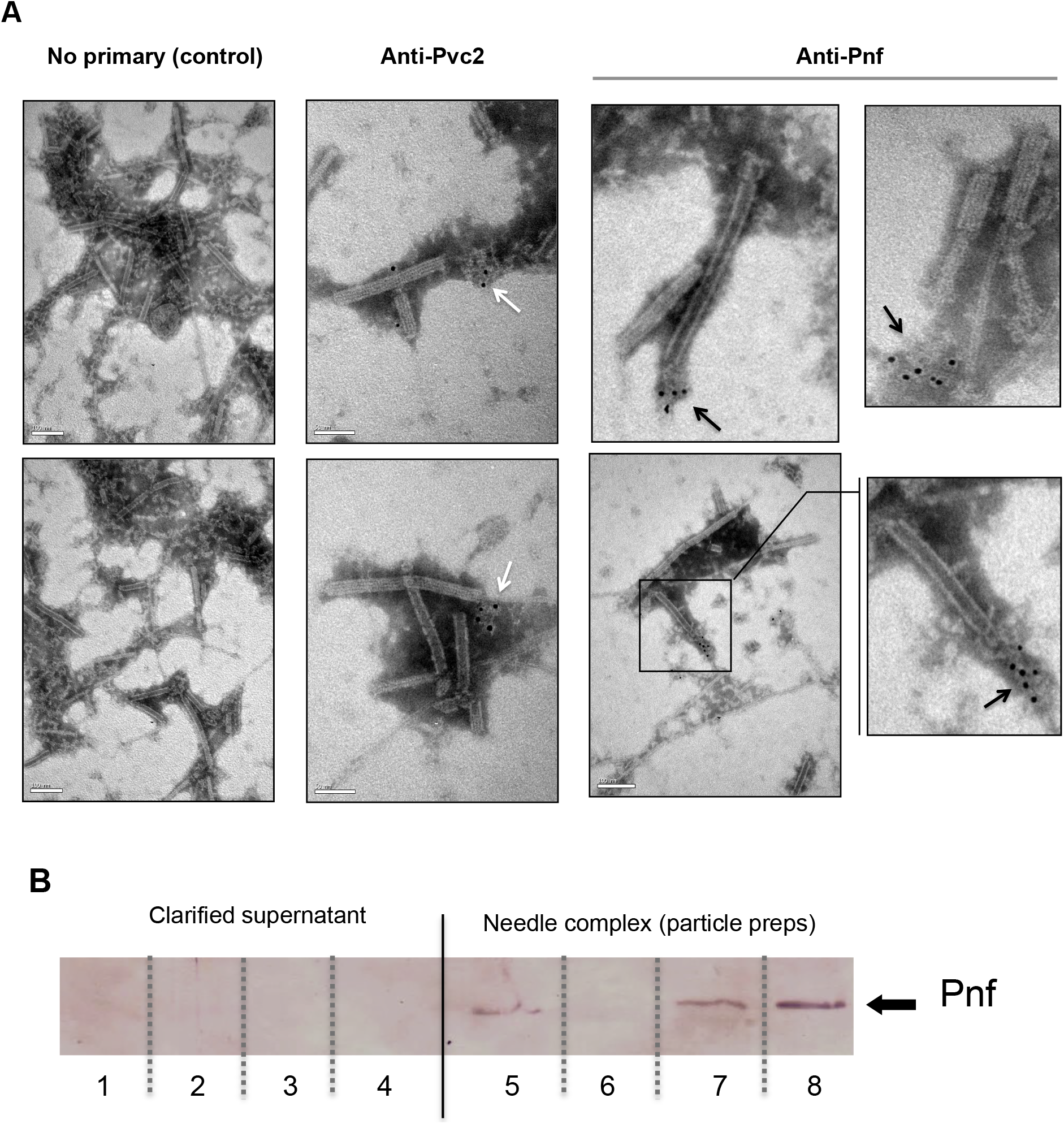
(**A**) Representative images of immuno-gold stained transmission electron microscopy grids confirming the Pnf-payload toxin is associated with the needle complex. PVC*pnf* needle complexes (PVC-NC) were prepared from supernatants of the *E. coli* 4df10 cosmid clone, which encodes the PVC*pnf* operon. We used anti-peptide antibodies against Pvc2 (DGPETELTINGAEE) and Pnf (TGQKPGNNEWKTGR) epitopes to localise these protein subunits. The Pvc2 epitope appeared to only become accessible to the antibody when subunits were “broken off” the ends (white arrows). The Pnf toxin could also only be detected at the ends of broken or contracted suggesting they are contained within the complex (black arrows). **(B)** Western blot analysis confirms that the Pnf protein can only be detected using the anti-peptide antibody if the needle complex is either chemically or physically disrupted. These preparations were taken from *Pa*^ATCC43949^ supernatants. The inability to detect Pnf in clarified supernatants confirms all the protein is associated with the PVC-NC enrichment preparation. Lanes 1+5; sonicated samples, 2+6; 1M NaCl treatments, 3+7; 1% SDS treatments 4+8; 1M Urea treatments. Note the PVC-NC appears stable in 1M NaCl.

Secondly we enriched needle complexes from insect toxic supernatants of an *E. coli* cosmid clone that encodes the *Pa*^ATCC43949^ PVC*pnf* operon, as previously described [31]. The anti-Pnf antibody was used for *in situ* labelling of Pnf on Transmission Electron Microscopy grids, visualised with negative staining and an anti-rabbit gold-conjugate secondary antibody. It was only possible to detect Pnf protein near the ends of either contracted or damaged needle complexes (Figure 3A). Note we saw no non-specific labelling when the gold-conjugate secondary antibody was used alone. In the case of the Pvc2 antibody, we only saw a signal associated with what appeared to be disrupted fragments of needle complexes suggesting the Pvc2 epitope is not normally solvent exposed in intact needle complexes.

### The Pnf protein requires delivery into the eukaryotic cell cytoplasm to exert its effect

In a previous publication we reported that injection of an enriched *Pa*^ATCC43949^ PVC*pnf* needle complex preparation; heterologously produced by an *E. coli* cosmid clone, caused melanisation and death of *Galleria mellonella* larvae within 30 minutes. In addition, microscopic analysis of phalloidin stained hemocytes taken from these dying animals revealed the cells were shrunken with highly condensed cytoskeletons, and likely already dead. This effect was abolished by heat denaturing the preparation. In this same publication [31] we demonstrated that transient cytoplasmic expression of the Pnf protein caused extensive cytoskeleton re-arrangement and likely cell death in cultured human HeLa cells, similar to that observed in the *ex vivo G. mellonella* hemocytes. In an attempt to directly visualise the interaction of the heterologously produced PVC*pnf* needle complex with insect hemocytes and to determine the initial effects on the cellular morphology, we injected intact or heat denatured PVC*pnf* needle complex preparations into 5^th^ instar *Manduca sexta* larvae before bleeding the animals and preparing their circulating hemocytes for surface examination by cryo-SEM. The surface of hemocytes injected with intact complex showed membrane ruffling consistent with the predicted mode of action of the Pnf protein (see below). Furthermore, we could also see linear structures approximately 150nm in length on the surface of the cells near the sites of membrane ruffles consistent with attached needle complexes. The surface of the control hemocytes injected with heat-denatured complex remained relatively smooth and homogeneous and we saw no equivalent linear structures. Figure 4 shows representative images from these experiments.

**Figure 4.**
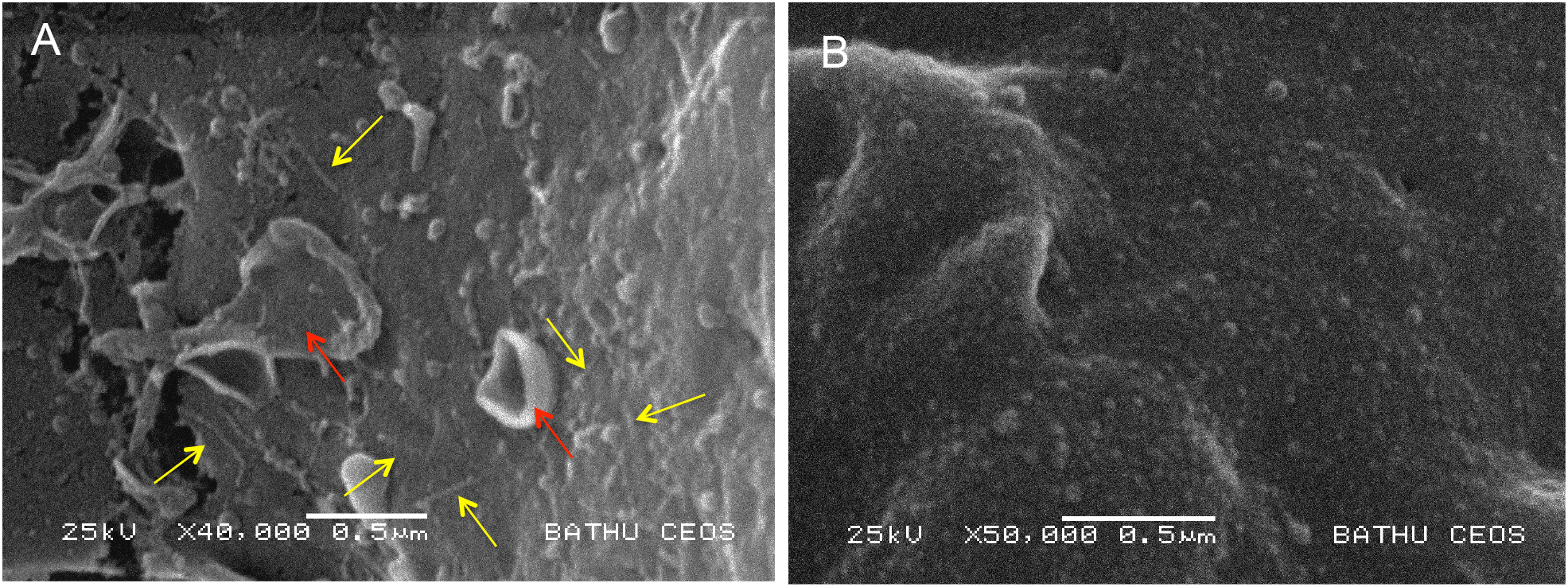
Cryo-SEM analysis of *ex vivo* hemocytes from 5^th^ instar *Manduca sexta* that had been injected with a native (A) or heat inactivated (B) enriched preparation of *Pa*^ATCC43949^ PVC*pnf* needle complexes heterologously produced by the *E. coli* cosmid clone. Note the abundant linear structures believed to be the PVC needle complex (yellow arrows) and membrane ruffling effect (red arrows) absent from the control treatment.

We wished to know if the Pnf effector could exert this toxic effect independently of the needle complex, when applied externally to eukaryotic cells. Therefore we heterologously expressed (in *E. coli*) and purified the Pnf protein in addition to a predicted toxoid derivative. The toxoid was designed based on homology between Pnf and the CNF2 toxin active site, wherein we mutated the cysteine at amino acid position 190 into an alanine (Pnf ^C190A^). Firstly, neither purified wild type nor toxoid proteins had any obvious toxic effect when injected into cohorts of *G. mellonella*, even at high doses (data not shown). We subsequently used bioPORTER, a liposome based transfection system, to introduce the purified proteins directly into cultured human cells. We visualised effects on the cytoskeleton and nucleus using TRITC-phalloidin and DAPI staining respectively. The wild type Pnf protein had a very clear effect on the cells, producing phenotypes consistent with those predicted by similarity to the CNF2 toxin. CNF2 is known to modify the cellular Rho GTPases, RhoA, Rac1 and Cdc42. Pnf delivery as a bioPORTER formulation lead to the formation of F-actin filaments within 24h followed by multi-nucleation by 48h, phenotypes consistent with the modification of the Rho GTPases. The toxoid derivative, delivered at the same dose using the same approach, produced no changes, giving cellular phenotypes consistent with that of the negative control or of the wild-type Pnf protein topically applied without the bioPORTER transfection agent (Figure 5).

**Figure 5.**
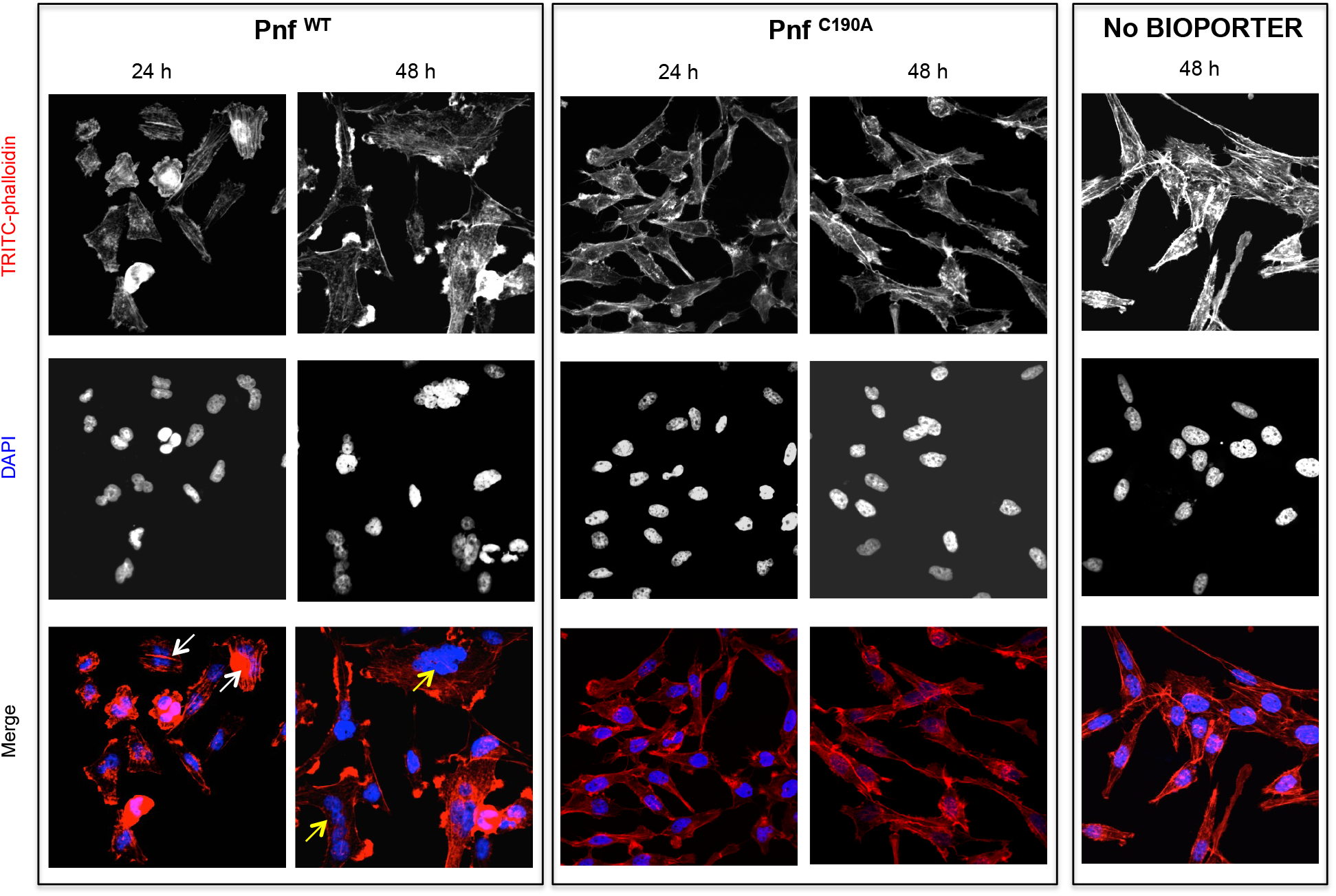
Pnf needs to gain access to the host cell cytoplasm to induce F-actin formation and multi-nucleation in HeLa cells. Wild-type and inactive toxoid mutant Pnf protein was delivered topically using BioPORTER. Cell cytoskeleton is stained with TRITC-Phalloidin and the cell nuclei with DAPI. This gave rise to phenotypes consistent with the molecular targets and that of the *Yersinia* CNF2 protein homologue. Note we see the F-actin formation by 24 h (white arrows) preceding extensive multi-nucleation of the host cell by 48 h (yellow arrows). Note neither application of the Pnf toxoid in a BioPORTER formulation or the purified wild-type Pnf protein without BioPORTER had any observable effect on the cells.

### The Pnf protein effector modifies small Rho GTPases

Based on homology to CNF2 the effect of Pnf on target cell proteins is predicted to include the modification of several Rho-family GTPases. Therefore we used western blot assays to examine *in vitro* transglutamination and deamidation effects of purified heterologously produced Pnf on purified small GTPases RhoA, Rac1 and Cdc42. Transglutamination is the formation of a covalent bond between a free amine group, as may be found on a lysine residue, and the gamma-carboxamide group of glutamine. As a result protein electrophoretic mobility of the protein is altered. Deamidation is a chemical reaction in which an amide functional group is removed from the protein, which may be detected using deamidated protein specific antibodies. These experiments demonstrated that Pnf induced transglutamination and deamidation of both RhoA and Rac1 (Figure 6), although unlike the reported activity of CNF2, had no effect on Cdc42. As predicted the active site toxoid mutant had no enzymic activity on any of the three Rho GTPases confirming it was a true toxoid derivative.

**Figure 6.**
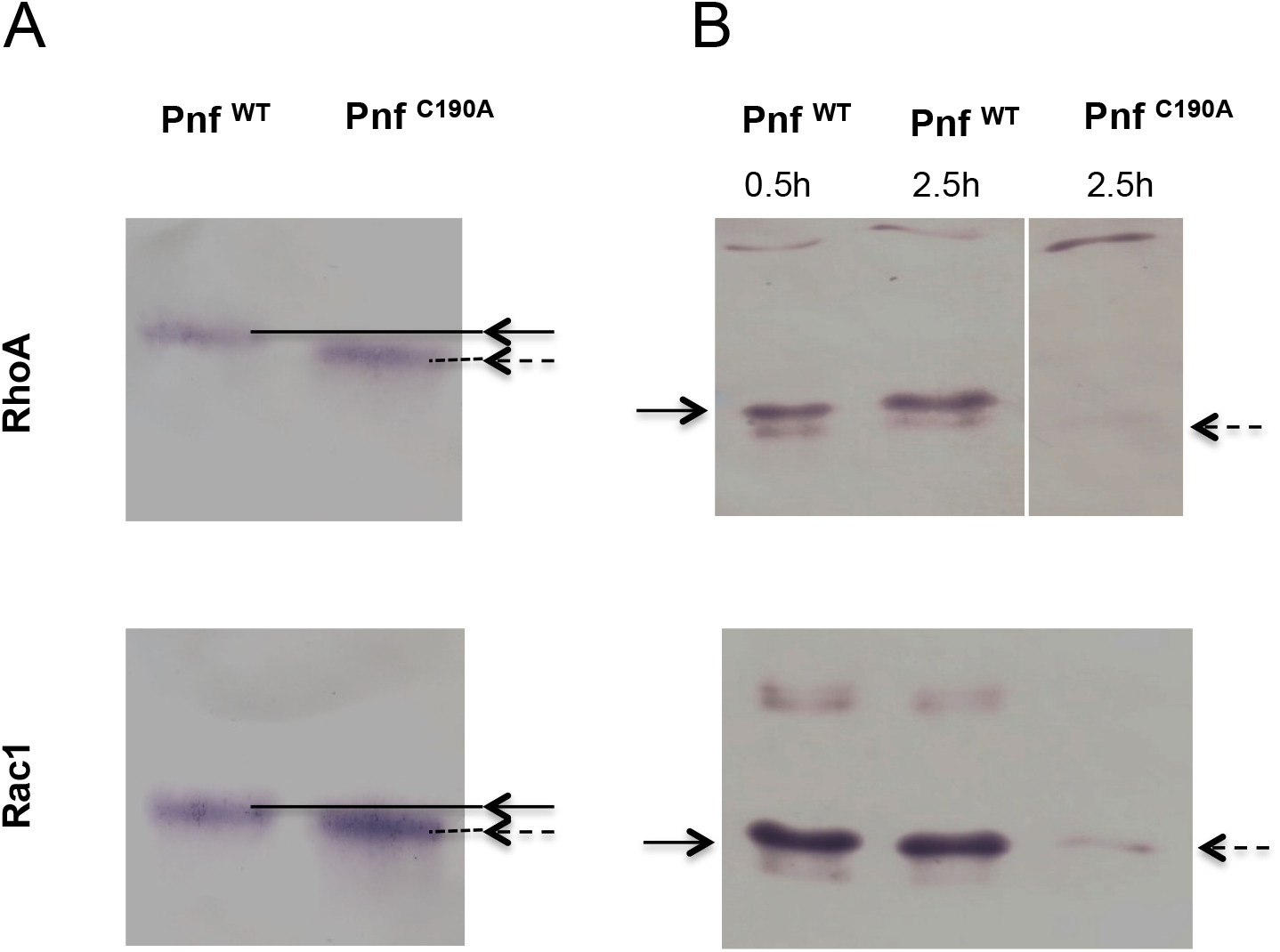
Pnf transglutaminates and deamidates purified mammalian RhoGTPAses at Gln63 (RhoA) and Gln 61 (Rac1). (**A**) For transglutination assays a 2:1 molar ratio of small Rho GTPase to purified Pnf was incubated in transglutination buffer in the presence of ethylediamine for 1 hour at 37°C. Note transglutinated GTPase runs slightly slower on the gel as visualised using anti RhoA and Rac1 antibodies. (**B**) For deamidation assays a 20:1 molar ratio of Rho GTPases; RhoA and Rac1, to purified Pnf toxin was incubated in deamidation buffer for either 30 min or 2.5 hours at 37 °C. Note deamidation is detected using an antibody specific towards deamidated Rho GTPase antigens. In both cases, the wild-type Pnf protein was active (solid arrows) while a site directed amino acid C190A toxoid mutant (in the predicted Pnf active site) showed no activity (dotted arrows).

## DISCUSSION

An analysis of the different subunit proteins of PVCs show they share several elements in common with other contractile phage-tail derived systems, including the Type VI secretion system (T6SS) [46] and to a lesser extent R-type pyocins [47]. However PVC-like elements are distinct in two important ways. Firstly, unlike the T6SS, they are freely released from the producing bacterial cell and so, in common with R-type pyocins, they can act at a distance. Secondly, like T6SS but unlike R-type Pyocins, they are evolved to inject bioactive protein effectors into other cells. We hypothesise that the PVCs are evolved to specifically target eukaryotic cells, unlike T6SS, which have been shown to be able to deliver to both eukaryotes and prokaryotic competitors. However, while our previous attempts to show PVC*pnf* attachment to a range of bacterial species from different genera showed no binding we could detect (data not shown), we cannot rule out the possibility that homologues exist which are able to target prokaryotes.

We speculate that these large protein complexes are costly for the cell to produce, consistent with the observation of population heterogeneity of *pvc*-operon expression. Indeed, uncontrolled heterologous over expression in *E. coli*, as cosmid clones, results in deletion of regions of the *pvc*-operon and loss of viability (unpublished data). It should be noted that in a natural insect infection the vast majority of the *Photorhabdus* bacterial population are sacrificial. The majority of the population act as a food source for the replicating nematodes, with very few cells passing into the next generation of infective juvenile nematodes [2]. As such the population may restrict PVC production to a limited number of sacrificial cells. The method of PVC release remains unclear, although to date we have not observed cell lysis associated with *pvc* expression. The finding that *Pa*^ATCC43949^ PVC*pnf*, and seven other *pvc*-operons from *Pa*^PB68^ and *Pl*^TT01^, all show population heterogeneity in expression, at least *in vitro*, suggests that they are likely deployed in a highly regulated and conservative manner. While it is difficult to fully characterise this heterogeneity *in vivo*, the PVC*pnf* GFP reporter strain did show restricted expression to one specific tissue, the spiracles, and not throughout the body of the insect. In regards to these experiments, it should be noted that we did not see any melanisation response around the bacterial biomass showing GFP. The insect melanisation immune response is typically activated at sites of encapsulation. This is mediated by the recruitment of hemocytes, surrounding and enclosing foreign bodies, and entombing them in melanin. The absence of melanisation around this GFP expressing bacterial mass is consistent with the expression of anti-hemocyte virulence genes, which are likely to include the native *Pl*^TT01^ *pvc*-operons.

Examination of the promoter regions has provided no clue as to the mechanism of population heterogeneity of expression. Nevertheless the identification of RfaH and a second cryptic conserved potential operator sequence upstream of the *pvc1* genes provides a starting point for addressing this in future. RfaH is a conserved anti-termination protein that is known to regulate large operons encoding for extracellular factors in *E. coli*. It is also believed to be important in ensuring appropriate transcriptional control of horizontally acquired operons [40]. Many of the *pvc*-operons in *Photorhabdus* and members of other genera (including *Xenorhabdus* and *Yersinia*) encode this operator sequence. An unusual example is the pADAP plasmid encoded *Serratia entomophila* anti feeding prophage (*afp*) [34]. While the *afp* promoter also encodes an RfaH operator sequence, it has been demonstrated that it is positively regulated by a tightly linked specific regulator protein, AnfA1 [48, 49]. This protein is a distant homologue of RfaH suggesting that other class I or III *pvc* operons are not necessarily under the regulation of the chromosomal RfaH orthologue, but might also be controlled by other diverse regulators that utilise this same operator sequence [50]. Operons containing the second cryptic putative regulatory sequence include [*Pl* TT01PVC*lopT*] and [*Pl* ^TT01^ PVC*u4*]. Analysis of the supplementary data from a recently published RNA-seq study [51], suggests that these operons may be dependent upon Hfq/HexA activity [52].

Unlike many of the other genera in which we see *pvc*-like operons, *Photorhabdus* genomes encode multiple copies, typically around 5 to 6, suggesting they play important and diverse roles in the life cycle. With this in mind, we examined the conservation of the different subunit genes between operons. We observe a “break point” in conservation, toward the 3’ end of the operons. We postulate this may be due to imprecise recombination events in the 3’ payload regions of *pvc*-operons, where incoming sequences, which have a GC-content that is distinct from the host genome, gradually ‘erode’ the upstream sequence. Alternatively, it is plausible that the lower GC at the distal end of these long operons (each of ∼25kb) may assist in strand separation during transcription, maintaining stoichiometry for these large, multi-subunit structures. Indeed low GC stretches of DNA are common origins of replication because of their reduced strand separation energy [53]. However, as yet we do not know whether the *pvc1* promoter serves the whole operon, or if there are additional promoters internal to the operon.

Each of the *pvc*-operons in a *Photorhabdus* genome encodes multiple paralogous copies of *pvc1/5* and *pvc2/3/4* genes. We were therefore surprised not to see any operons showing signs of genetic degradation. This suggests there is sufficient positive selection for maintaining these multiple operons, with each operon potentially adapted for a specific role. This hypothesis is supported by the high variation in the Pvc13 protein sequences, which we speculate represent the host cell binding fibres. The need to maintain multiple copies of *pvc*-operons may also have arisen if the structural genes for the needle complexes are specifically adapted for delivery of their cognate *cis*-linked effector proteins in some way.

Circumstantial evidence from genomic sequences and previous work on the related AFP system of *Serratia* has suggested the needle complexes serve to deliver the *cis*-encoded effector proteins. We present here for the first time direct evidence that a linked effector protein does in fact become physically associated with the needle complex. Western blot detection of Pnf from preparations enriched for needle complexes taken from the native *Pa*^ATCC43949^ supernatants confirmed it was being expressed *in vitro* and suggested it was physically associated with the complexes. In addition, physical or chemical disruption was required to release the Pnf protein for detection. When taken alongside the immuno-gold EM observations, showing Pnf could only be seen near contracted or damaged needle complexes, it confirms the protein is either sequestered inside the complex or physically associated in such as way that the TGQKPGNNEWKTGR epitope is not solvent accessible. The anti-Pvc2 antibody is able to specifically detect the protein in Western blots, however it only showed binding to what appeared to be disrupted fragments of needle complexes, again suggesting the relevant epitope is not accessible in the intact native needle complex structure. Indeed, iTasser structural model simulations of a PVC outer sheath Pvc2 protein, using the homologous *Pseudomonas* 3J9Q PDB structure of an R-type pyocin outer sheath as a model [54], supports this idea, suggesting the epitope is partially occluded between adjacent subunits.

While we have not yet directly demonstrated injection of Pnf into host cells by the needle complex, the results of the topical application and bioPORTER transfection experiments confirmed that the Pnf effector absolutely requires a mechanism to facilitate entry into the host cell cytoplasm to exert its effect. We argue the evidence for injection by the needle complex is very strong, and is corroborated by the SEM visualisation of needle-like structures of the correct dimensions on the surface of intoxicated hemocytes. Finally, we have confirmed that Pnf acts in a manner similar to the *Yersinia* CNF2 toxin, modifying two of the same Rho-GTPases, which correlates with the observed phenotypic effects on the cell.

## MATERIALS AND METHODS

### Insects, bacterial strains and growth conditions

*Manduca sexta* (Lepidoptera: *Sphingidae*) were individually reared as described [55]. Briefly, larvae were maintained individually at 25°C under a photoperiod of 17 h light: 7 h dark and fed on an artificial diet based on wheat germ. Larvae 1 day after ecdysis to the 5th instar were used for all experiments. Batches of wax moth larvae (75 g; Livefood UK Ltd, Rooks Bridge, UK) in their final instar stage were stored in the dark at 4°C and used within a week of receipt. DH5α™ *E. coli* (containing various plasmid constructs) were grown on LB agar at 37°C or in LB liquid, shaking at 200 rpm. Spontaneous rifampicin-resistant mutants of *Photorhabdus asymbiotica subsp. asymbiotica* Thai (strain PB68.1) [56] and *Photorhabdus luminescens subsp. laumondii* TTO1 [12] were used in these studies as hosts for reporter plasmids. *Photorhabdus* were routinely cultured in LB broth or on LB agar supplemented with 0.1 % (w/v) pyruvate at 30°C or 37°C (for *P. asymbiotica*). When required antibiotics were added at the following concentrations: ampicillin (Amp): 100 μg ml^-1^, kanamycin (Km): 25 μg ml^-1^, chloramphenicol (Cm): 25 μg ml^-1^, rifampicin (Rif): 25 μg ml^-1^. HeLa ATCC CCL2 cells were cultured for 10 passages in Dulbecco’s modified Eagle medium (Sigma-Aldrich) containing 4.5 g/L glucose (Sigma-Aldrich), 10% heat-inactivated fetal bovine serum (Sigma-Aldrich), 2 mM glutamine (Sigma-Aldrich), 100 μg /mL penicillin, and 100 μg/mL streptomycin (Sigma-Aldrich) and incubated at 37°C and 5% CO_2_.

### PVC gene reporter plasmid construction

Translational fusions with the *gfpmut*2 gene were constructed by PCR in a pACYC184 vector containing the *gfpmut*2 (pACYC-GFP) [57] as follows. The *pvc1, pnf and rpsM* genes (consisting of promoter regions and the first 150 bp of coding sequence) were amplified from *P. asymbiotica* ^ATCC43949^ genomic DNA and cloned into pACYC-*gfp* to generate pACYC-*afp1*-*gfp*, pACYC-*pnf*-*gfp* and pACYC-*rpsM*-*gfp*. The constructs were further digested to release the *pvc1, pnf or rpsM* genes in frame with *gfp* and the fusion fragments were cloned into pBBR1-MCS [58] to generate pBBR1-*pvc1*-*gfp*, pBBR1-*pnf*-*gfp* and pBBR1-*rpsM*-*gfp*. Mating experiments were performed as previously described [59] to transfer plasmid constructs into *P. luminescens* ^TTO1^ resulting in *Pl*^TTO1^*-pvc1*-*gfp, Pl*^TTO1^*-pnf*-*gfp* and *Pl*^TTO1^*-rpsM*-*gfp*. Plasmid stability was confirmed in bacteria harbouring the various constructs isolated after *in vivo* passages. For the expanded panel of *gfp*-reporter fusions, the promoter regions for the operons selected, inclusive of the putative RfaH operator sites, and the native RBS and first codon of the *pvc1* gene (approximately 500 bp upstream), were cloned in to the pAGAG vector. pAGAG was derivatised from the promoterless pGAG1 *gfp* bearing plasmid., In brief, pGAG1 was used as a template to amplify *gfpmut3\** without a start codon using primers pG_GFPfor (5’-AATGTCGACCGTAAAGGAGAAGAACTTTTC-3’) and pG_GFPrev (5’-AATACTAGTGGATCTATTTGTATAGTTCATCCATG-3’). The resulting product and the pGAG1 vector were cut by digestion with SalI-HF and SpeI and ligated together, thus replacing the original intact *gfpmut3\** gene with one that lacks a ribosome binding site and the first ATG codon. Thus, 5’ regions introduced subsequently restored the construct. All upstream regions were incorporated between the KpnI and BamHI sites of the resulting pAGAG vector. The Pnf reporter discussed in this paper specifically (Figure S4), was amplified using the primers (PB68.1Pnf-BamHI F 5’-ATA<u>GGATCC</u>ATCCCAACGTATCTTGTCC-3’ and PB68.1Pnf-KpnI R 5’-ATTGGTACCTGTACTTGTAGACATAAAAGCCC-3’

### Fluorescent reporter strain assays

#### *In vitro* experiments

Reporter strains were cultured with shaking aeration in LB liquid medium supplemented with 20% (v/v) freshly harvested 5^th^ instar *M. sexta* clarified hemolymph. To obtain the hemolymph, insects were chilled on ice for 20 minutes before being bled (by cutting the tip of the tail horn) into a tube on ice, containing 10 μl of saturated Phenol Thio Urea (PTU) solution, which prevents melanisation. Hemolymph was clarified by centrifugation to remove hemocytes and other debris. Bacteria were grown to late stationary phase, before microscopic visualisation using a Leica inverted epi-fluorescent microscope. ***In vivo* experiments**, we injected ca. 100 cells of the reporter strains into cohorts of 5^th^ instar *M. sexta*, and allowed the infection to establish before macroscopic examination of insect tissues using a (fluorescence) dissecting microscope. We also took hemolymph samples from these insects and performed microscopic examination of fixed *ex vivo* hemocytes stained with phalloidin conjugate and confocal microscopy to visualise host cell cytoskeleton and any GFP expression from the recombinant bacteria. Images were acquired with a LSM510 confocal microscope (Leica).

### PVC purification from *E. coli* cosmid clone supernatants and electron microscopy

Cosmid libraries of *P. asymbiotica* ^ATCC43949^ were prepared in *E. coli* EC100 and arrayed into 96-well microtiter plates by MWG Biotech, Munich, Germany, as described previously [13, 28]. A 250ml overnight culture of *E. coli* with the *Pa*^ATCC43949^ PVC*pnf* cosmid (c4DF10) was grown in LB medium supplemented with 100 μg ml^-1^ ampicillin at 28°C with aeration in the dark. The culture was centrifuged at 6800 x *g* at 4°C for 30 min at 4°C. The supernatant was decanted to remove each cell pellet, and the centrifugation procedure was repeated to remove any remaining cells. Cell-free supernatants were then centrifuged, in small batches, at 150,000 × *g* for 90 min at 4°C to harvest particulate material. The particulate pellets were washed by gentle re-suspension in 1× Phosphate Buffered Saline (PBS) before a second centrifugation at 150,000 × *g* for 90 min at 4°C to pellet the particulate material. Each pellet was further separated by DEAE-Sepharose chromatography. 10 ml of particulate material in ice-cold PBS were mixed with an equivalent volume of DEAE-Sepharose CL-6B anion exchanger (in PBS) and the preparation was incubated at room temperature for 15 min. The Sepharose resin was harvested by centrifugation (3,000 × *g*), and the supernatant was discarded. The resin was resuspended in 40 ml of ice-cold PBS and again harvested by centrifugation. This washing step was repeated another three times, and the resin was finally resuspended in 10 ml of elution buffer (0.5 M NaCl, 50 mM phosphate buffer [pH 7.4]). The resin was removed by centrifugation, and the supernatant containing the PVCs was again centrifuged at 150,000 × *g* for 90 min at 4°C to pellet the particulate material and concentrate the needle structures in 500 μl of ice-cold PBS.

For transmission electron microscopy (TEM) pioloform-covered 300-mesh copper grids that were coated with a fine layer of carbon were used as substrates for the protein fractions. The following four aqueous negative stains were tested with the protein samples: 1% uranyl acetate, 3% ammonium molybdate, 3% methylamine tungstate, and 2% sodium silicotungstate. The preferred stain, 3% methylamine tungstate, produced acceptable contrast and minimum artefacts and was subsequently used for all samples viewed by TEM. The coated grids were exposed to UV light for 16 h immediately prior to use to ensure adequate wetting of the substrate. A 10 μl drop was applied to the TEM grid, and the protein was allowed to settle for 5 min. Liquid was absorbed with filter paper from the edge of the grid and replaced immediately with 10 μl of filtered negative stain. The drop was partially removed with filter paper, and the grids were allowed to air dry thoroughly before they were viewed with a JEOL 1200EX transmission electron microscope (JEOL, Tokyo, Japan) operating at 80 kV.

### Pnf cloning and heterologous expression for *Galleria* injection and antibody specificity test

*Pnf* gene was amplified from *P. asymbiotica* ^ATCC43949^ genomic DNA (using primers Pnf_NdeI 5’-ATATAT<u>CATATG</u>ATGTTAAAATATGCTAATCCT-3’, Pnf_BamHI 5’-ATATAT<u>GGATCC</u>TTATAACAACCGTTTTTTAAG-3’) and the PCR product was purified and cloned in-frame with a His-tag into the IPTG-inducible expression plasmid pET-15b (Novagen) to create construct pET15b-Pnf. The clone was verified by sequencing and transformed into Arctic Express competent cells (Agilent) for protein expression. A site-directed mutant of Pnf (toxoid) was generated with either the QuikChange site-directed mutagenesis kit (Agilent). To construct the Pnf mutant plasmid pET15b-Pnf_C190A_, pET15b-Pnf was amplified with FPLC-purified primers designed to generate a Cys to Ala substitution at position 190 (Pnf_C190A__for 5’-TCACCGAATATACCATAGTAGCACCGCTCAATGCTCCAGAC-3’, Pnf_C190A__rev 5’-GTCTGGAGCATTGAGCGGTGCTACTATGGTATATTCGGTGA-3’) using the following thermal profile (step 1: 95°C for 30 s, step 2: 95°C for 30 s, 55°C for 60 s, 68°C for 6 min 45 s for 16 cycles). The identity of six different positive clones was confirmed by sequencing. Subsequently, −80°C glycerol stocks were used to inoculate 5 ml of fresh LB medium supplemented with 0.2% (w/v) glucose and 100 μg ml^-1^ ampicillin. Bacteria were grown overnight at 30°C with shaking, and 1 ml of the culture was then harvested, re-suspended in 100 ml of the same medium, and incubated in an orbital incubator at 37°C until the optical density at 600 nm was 0.7 to 0.9. Cells were then harvested at room temperature by centrifugation at 4,000 rpm for 10 min. The pellet was re-suspended in 100 ml of fresh LB medium supplemented with the 100 μg ml^-1^ ampicillin and 0.1 mM of the inducer isopropyl-β-d-thiogalactopyranoside (IPTG). Optimized times for inductions were determined experimentally, and cells were then harvested. The bacterial cell pellet was re-suspended in 10 ml of 1x PBS and sonicated (four 20-s sonications at 45 mA using a Branson 450 digital Sonifier) fitted with a tapered probe. The freshly sonicated samples were then diluted in 1x PBS for injection into *Galleria* larvae and for SDS-polyacrylamide gel electrophoresis analysis to confirm expression of the target protein. For toxicity testing cohorts of *Galleria* larvae (n=20) were chilled on ice before injection with 10 μl of a dilution series (in sterile PBS) of sonicated cells expressing Pnf or vector control. Insects were then returned to room temperature and observed for 5 days or mortality or morbidity.

### Recombinant Pnf and small Rho-GTPase purification

ArcticExpress containing pET-15b-Pnf were initially grown in LB broth supplemented with 100 μg ml^-1^ ampicillin at 37°C until OD 0.6 when Pnf expression was induced with a final concentration of 0.1 mM of IPTG at 12°C for 16 h to produce soluble Pnf. Pnf was purified over HisTrap™ Ni^2+^-affinity column with the fast phase liquid chromatography (FPLC) AKTA system as per the manufacturer’s protocol (GE Healthcare). Plasmids pGEX-2T-wtRhoA, pGEX-2T-wtRac1 and pGEX-2T-G25K (Cdc42) were gifts from Prof Alan Hall (University College London, London, UK) and were maintained in *E. coli* DH5α grown on LB agar or in LB broth supplemented with 50 μg ml^-1^ ampicillin. RhoA, Rac1 and Cdc42 were purified over GSTrap HP™ affinity columns with the FPLC AKTA system as per the manufacturer’s protocol (GE Healthcare).

### BioPORTER assay and actin stress fibre analysis

For BioPORTER assays, 80 μl of purified wild type and mutant Pnf proteins (500 μg ml^-1^), or PBS as a negative control, were added to one BioPORTER tube (Genlantis) and resuspended in 920 μl of DMEM. The samples were added to HeLa cells grown in 6-well plates and incubated for 4 h. BioPORTER/protein or PBS mixes were replaced by fresh complete medium and the cells were incubated for 20–48 h. To visualize cell morphology and actin cytoskeleton, cells were fixed for 15 min in 4% PBS-formaldehyde, permeabilized with 0.1% Triton X-100 and stained with Tetramethylrhodamine B isothiocyanate (TRITC)-phalloidin (Sigma) and DAPI dihydrochloride (Sigma). Images were acquired with a LSM510 confocal microscope (Leica).

### Deamidation and Transglutamination of Rho GTPases

#### Deamidation assay

Deamidation assays were done according to previously described procedures [60] with the following modifications. Briefly, a 20:1 molar ratio of GTPase (RhoA, Rac1 or Cdc42) to toxin was incubated in deamidation buffer (50 mM NaCl, 50 mM Tris-HCl pH 7.4, 5 mM MgCl_2_, 1 mM DTT, 1 mM phenylmethanesulphonyl fluoride) for either 30 min or 2.5 h at 37°C. Untreated RhoA served as a negative control. After toxin treatment, samples were concentrated by the addition of 10% trichloroacetic acid and stored overnight at 4°C. Precipitated proteins were pelleted, washed with acetone, air-dried and resuspended in 20 mM Tris-HCl pH 7.4. Samples were subjected to SDS-PAGE and analysed by Western blotting using either an anti-RhoA (1:1500, Santa Cruz Biotechnology), anti-Rac1 (1:5000, Upstate Biotechnology), or anti-Cdc42 (1:1000, Santa Cruz Biotechnology) monoclonal antibody or rabbit polyclonal antisera (1:2000) that had been raised against a peptide antigen specifically designed to detect modified/deamidated RhoA/Rac1/Cdc42 [61], provided by Prof A. D. O’Brien, Department of Microbiology and Immunology at Uniformed Services University, Maryland, USA). Reactive proteins were detected with either the HRP-conjugated goat anti-mouse IgG (Sigma) or donkey anti-rabbit IgG (1:3000, Sigma) followed by visualization with DAB (Sigma). ***Transglutamination assay***: Transglutamination assays were done as previously described [62] with several modifications. Briefly, a 2:1 molar ratio of RhoA to toxin was incubated in transglutamination buffer (50 mM Tris-HCl pH 7.4, 8 mM CaCl_2_, 5 mM MgCl_2_, 1 mM DTT, 1 mM EDTA) in the presence of ethylenediamine (50 mM, pH 9) for 10 min or 1 h at 37°C. As a negative control, RhoA was incubated with ethylenediamine but without toxin. Samples (0.25 μg RhoA/well) were subjected to SDS-PAGE and then processed for Western blot analyses as described above. Immunoblots were probed with a mouse anti-RhoA monoclonal antibody (1:1500, Santa Cruz Biotechnology) and reactive proteins visualized with DAB after incubation with the HRP-conjugated goat anti-mouse IgG secondary antibody.

## ACKNOWLEDGEMENTS

This work would not have been possible without the much appreciated funding by BBSRC grants BB/C008367/1 and BB/E021328/1, The Leverhulme Trust grant RPG-2015-194, EPSRC (MOAC) DTP EP/F500378/1, MRC DTP in Interdisciplinary Biomedical Research MR/N014294/1 and the Warwick University Medical School. We would also like to acknowledge Chris Apark for maintaining and supplying the *Manduca sexta* insects from the University or Bath colony.

## SUPPLIMENTARY INFORMATION

### SUPPLIMENTARY METHODS

#### A bioinformatic analysis of *pvc* structural operon sequences

DNA sequences for each of the 16 conserved structural loci were clustered syntenically (all *pvc1*s, all *pvc2*’s etc.). % GC content for each CDS in each syntenic position was calculated (up to 16 observations per locus), and plotted as a boxplot via ggplot2 (Figure S1A). The average GC content across the full operon, as well as for the whole genome, were plotted as intervals in the plot background to show the PVC loci %GC in contrast. The breakpoint was defined by use of the “cumSEG” package in R [63]. Amino acid similarity scores (Figure S1B) were generated by CLUSTAL Omega [64] multiple sequence alignment, using default parameters. Resulting pairwise alignment scores were plotted as boxplots using ggplot.

#### RNA purification and RT-PCR

For *in vitro* transcription analysis, overnight cultures of *P. asymbiotica* were sub-cultured into liquid LB medium and grown with aeration at 28°C or 37°C 200 rpm in the dark. Planktonic cultures were collected at 4, 8 and 24 h and mixed with a double volume of RNAlater (Ambion) and after 5 minute incubation, bacteria were harvested by centrifugation and the pellets stored at −80°C. For *in vivo* transcription analysis, overnight cultures of *P. asymbiotica* were extensively washed in PBS and diluted in Grace’s insect media (GIM) to achieve 1000 bacteria per 50 μl of culture. Each *M. sexta* larvae was injected with 50 μl of *P. asymbiotica* culture and they were placed in a humid temperature controlled room at 28°C. After 3h or 6 h of incubation, insects were bled in equal volume of GIM containing 20mM phenylthiocarbamide (PTC). The sample was initially fractionated into plasma and total hemocytes by centrifugation at 200 x *g* at 4°C for 5 min, and plasma was further centrifuged at 6800 x *g* at 4°C for 5 min to form a bacterial pellet. For each condition, total RNA was extracted using the RNeasy Mini Kit (Qiagen) and 2 μg total RNA was treated with TURBO DNA-free Kit (Ambion) and subjected to RT-PCR using the Qiagen OneStep RT-PCR kit. Each RT-PCR reaction performed in a volume of 50 μl (containing 100 ng template RNA, 1x QIAGEN OneStep RT-PCR buffer, 400 μM dNTPs, 0.6 μM gene specific primers, 5U RNase inhibitor and 2 μl of QIAGEN OneStep RT-PCR enzyme mix) for 28 cycles.

## SUPPLIMENTARY FIGURES

**Figure S1.**
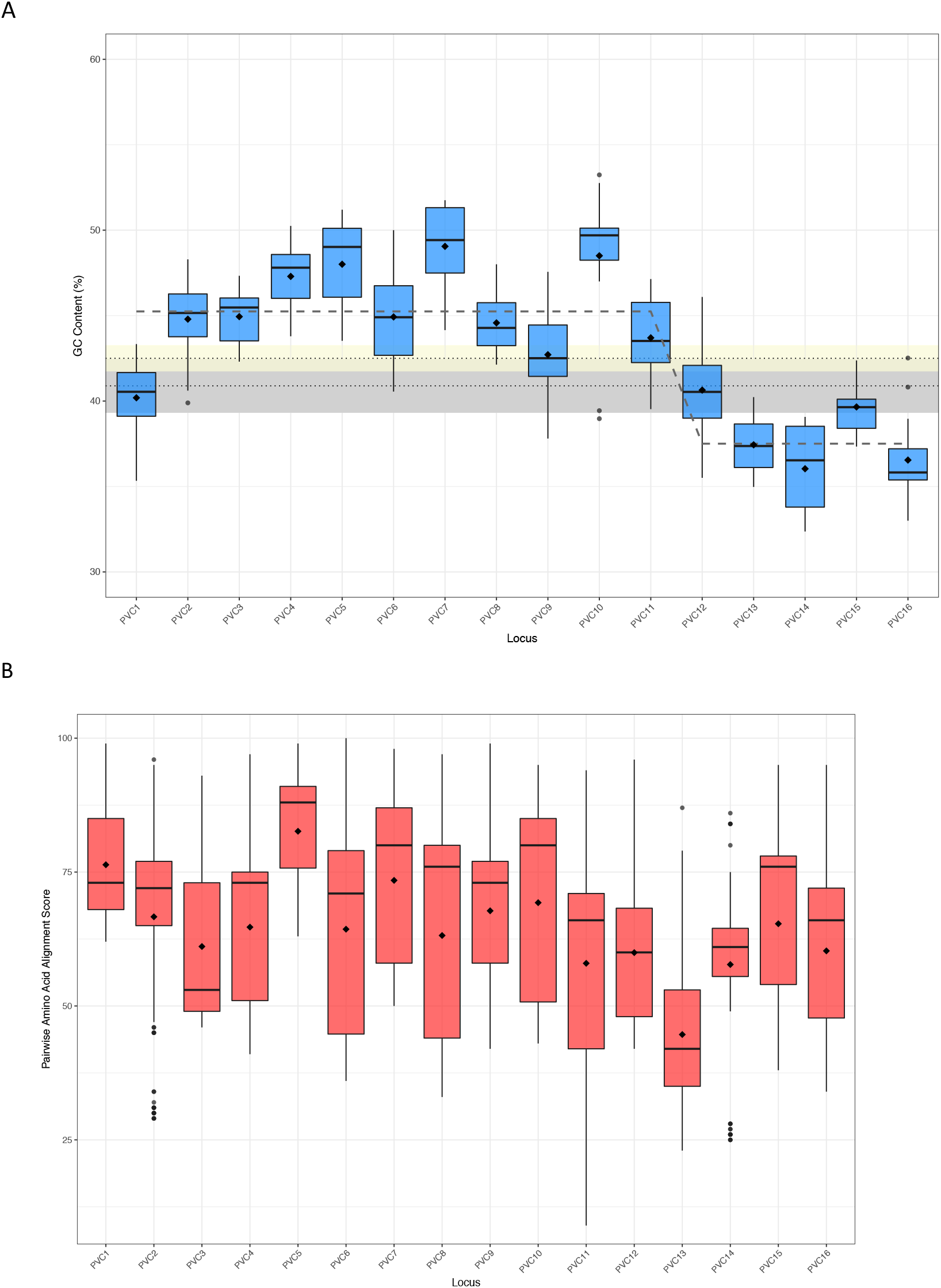
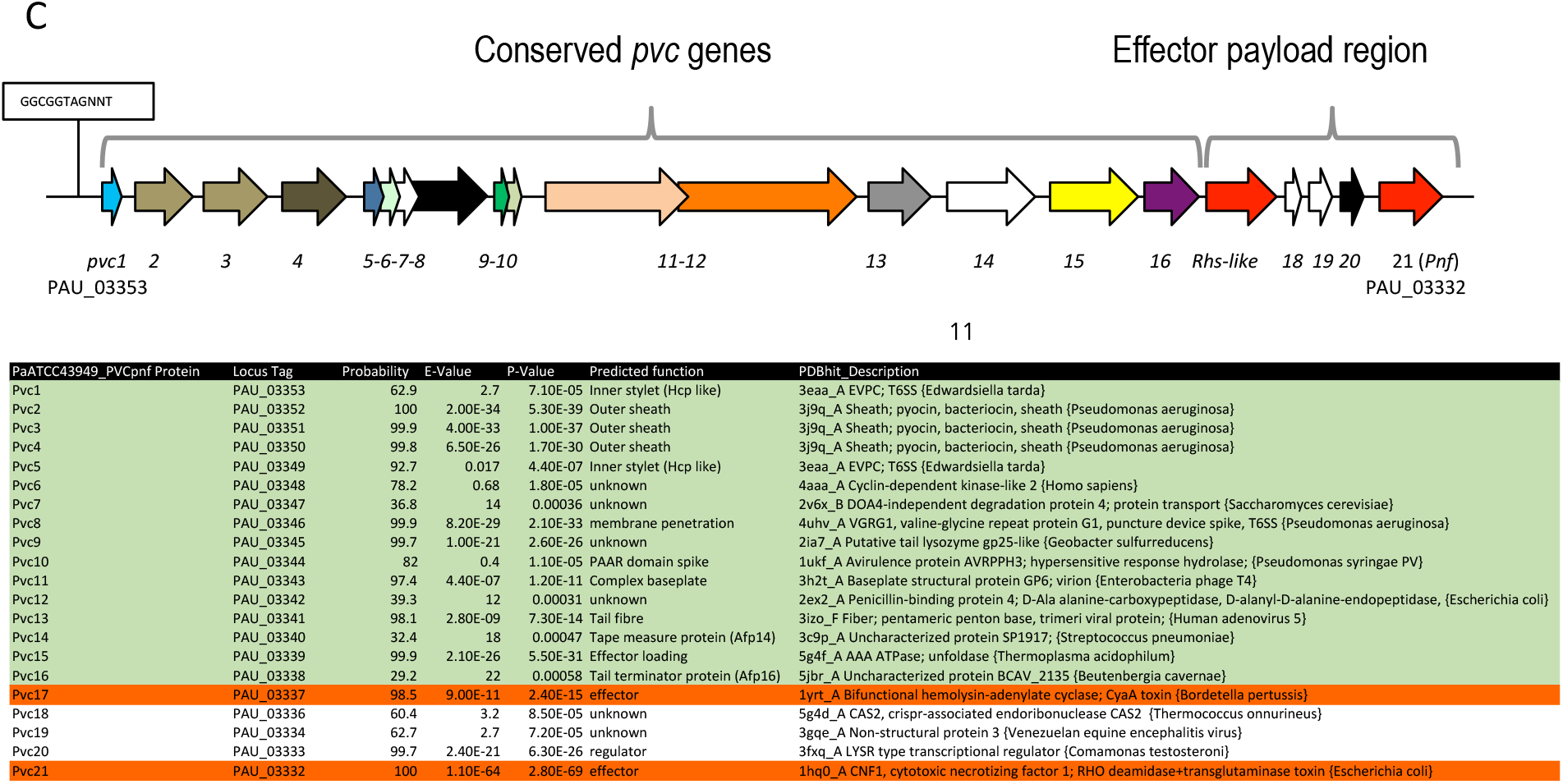
(**A**) Boxplots of the mean GC content across 16 different *pvc* operons of *Photorhabdus*. The GC was calculated for each of the 16 structural loci (clustered by annotated/predicted function and syntenic position in operon consistent with the nomenclature devised in this paper). GC content itself was calculated via a bespoke script, outputting data to be visualised in RStudio. Data was plotted and the step-function fit (black dashed line) was calculated using the mean GC value for each locus via the *cumSeg* package for breakpoint estimation in genomic sequences. Diamonds represent mean locus GC. Beige box shows the source genome mean (dotted line) GC content and standard deviation (upper and lower box bounds). Grey box shows the operon GC mean (dotted line) and standard deviation (upper and lower box bounds). (**B**) Box plots of amino acid similarity across homologous protein sequences for these same 16 operons. Amino acid sequences were clustered together as in (A), by annotation and syntenic position. Global Multiple Sequence Alignments were created with CLUSTAL Omega, using the default parameters (Gonnet transition matrix, gap open penalty 6 bits, gap extend 1 bit). Pairwise amino acid alignment scores were extracted from the CLUSTALO output and plotted in RStudio via bespoke scripts. Diamonds indicate mean pairwise alignment scores. Dots indicate pairwise values that are outliers, beyond 1.5 X the interquartile range (as automatically calculated by the ggplot2 package). (**C**) A map of the model class I *Pa*^ATCC43949^ PVC*pnf* operon showing two effector genes in the payload region in red/orange.

**Figure S2.**
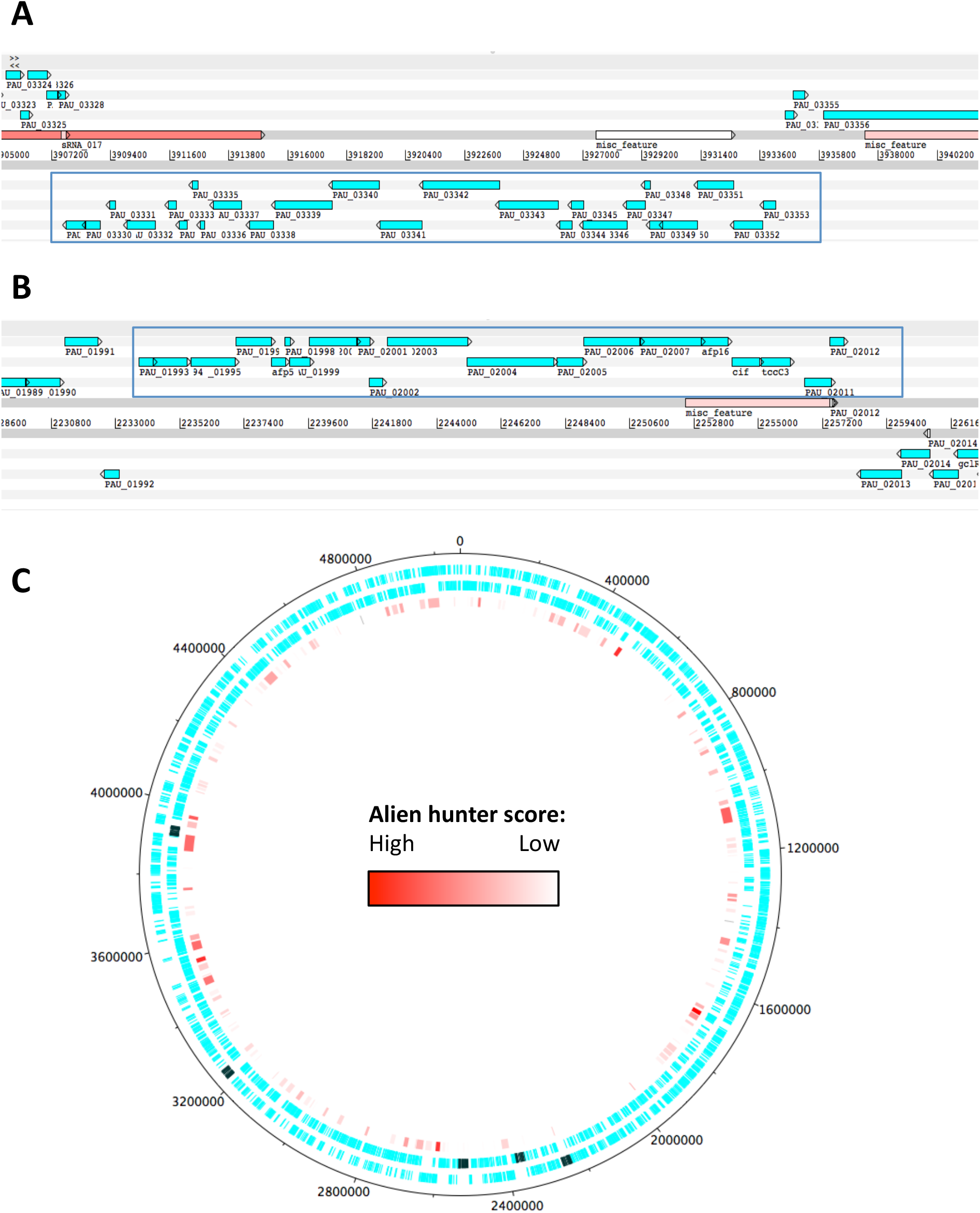
PVC operons in relation to putative regions of horizontal gene transfer as identified by Alien Hunter. (**A**) The PVC*pnf* and (**B**) the PVC*cif* operons are highlighted by the blue rectangle. Alien Hunter regions of HGT are designated by the features in tones of red. In red are the regions with the highest score and thus probability for HGT whilst in white are the regions with the lowest scores. **(C**) The *P. asymbiotica* ^ATCC43949^ chromosome. The first concentric circle denotes genes on the forward strand while the second circle denotes genes in the reverse strand. In dark green are the PVC operons. The third circle shows regions of HGT as identified by Alien Hunter.

**Figure S3.**
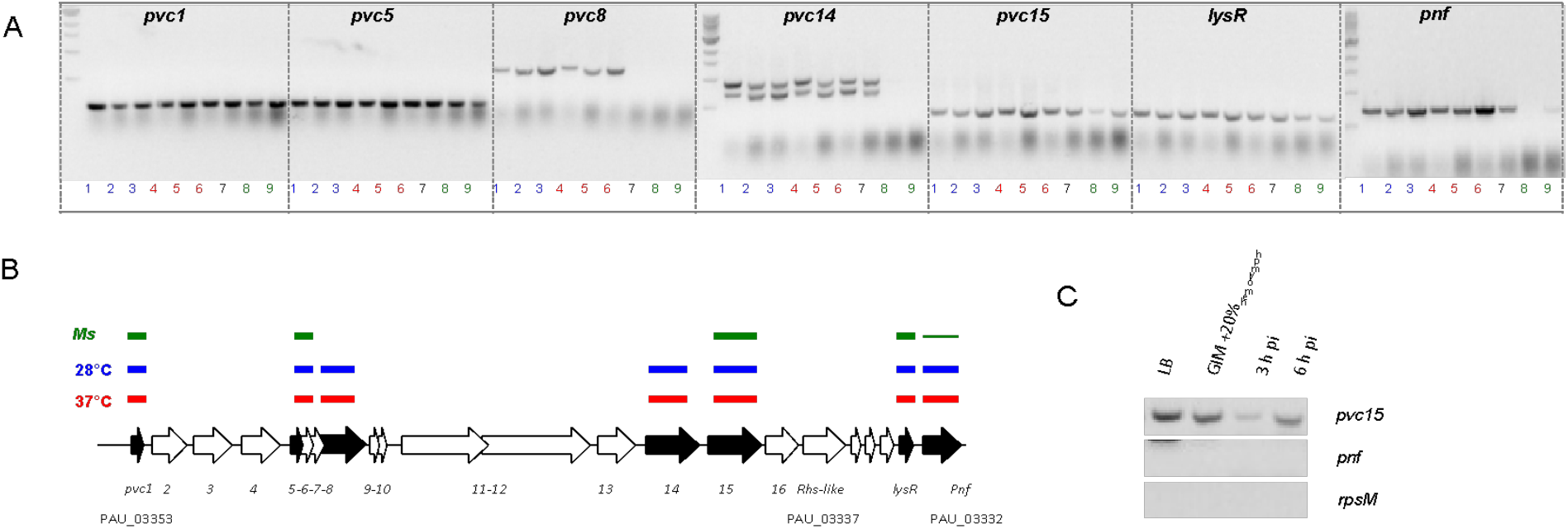
**(A)** RT-PCR analysis of gene transcription of various genes of the *Pa*^ATCC43949^ PVC*pnf* operon over time *in vitro* at insect (28°C) and human (37°C) relevant temperatures and *in vivo* during *Manduca sexta* (*Ms*) infection. Lane key; lanes 1, 2 and 3 (blue) represent *in vitro* growth in aerated LB at 28°C for 4, 8 and 24h respectively; lanes 4, 5 and 6 (red) are growth in aerated LB at 37°C for 4, 8 and 24h; lane 7 (black) is growth in LB at 28°C for 16h; lanes 8 and 9 (green) are from 3h and 6h post infection blood of *Ms* infected with *P. asymbiotica* at 28°C. **(B)** Map of the operon showing RT-PCR target genes in black. The lane-colour coded bars above the ORFs summarise in which conditions gene transcription could be detected. Note *pvc8* and *pvc14* mRNA could not be detected from infected *Ms* and the *pnf* mRNA was only detected after 6h of infection. **(C)** RT-PCR signals for *pvc15* and *pnf* from infected insects with the *rpsM* (ribosomal subunit protein S13) loading control. Lanes represent (in order); 4h growth in LB at 28°C; 4h growth in Grace’s insect medium supplemented with 20% (v/v) Ms hemolymph; 3h and 6h post infection *ex vivo* blood of *Ms* infected with *P. asymbiotica* at 28°C.

**Figure S4.**
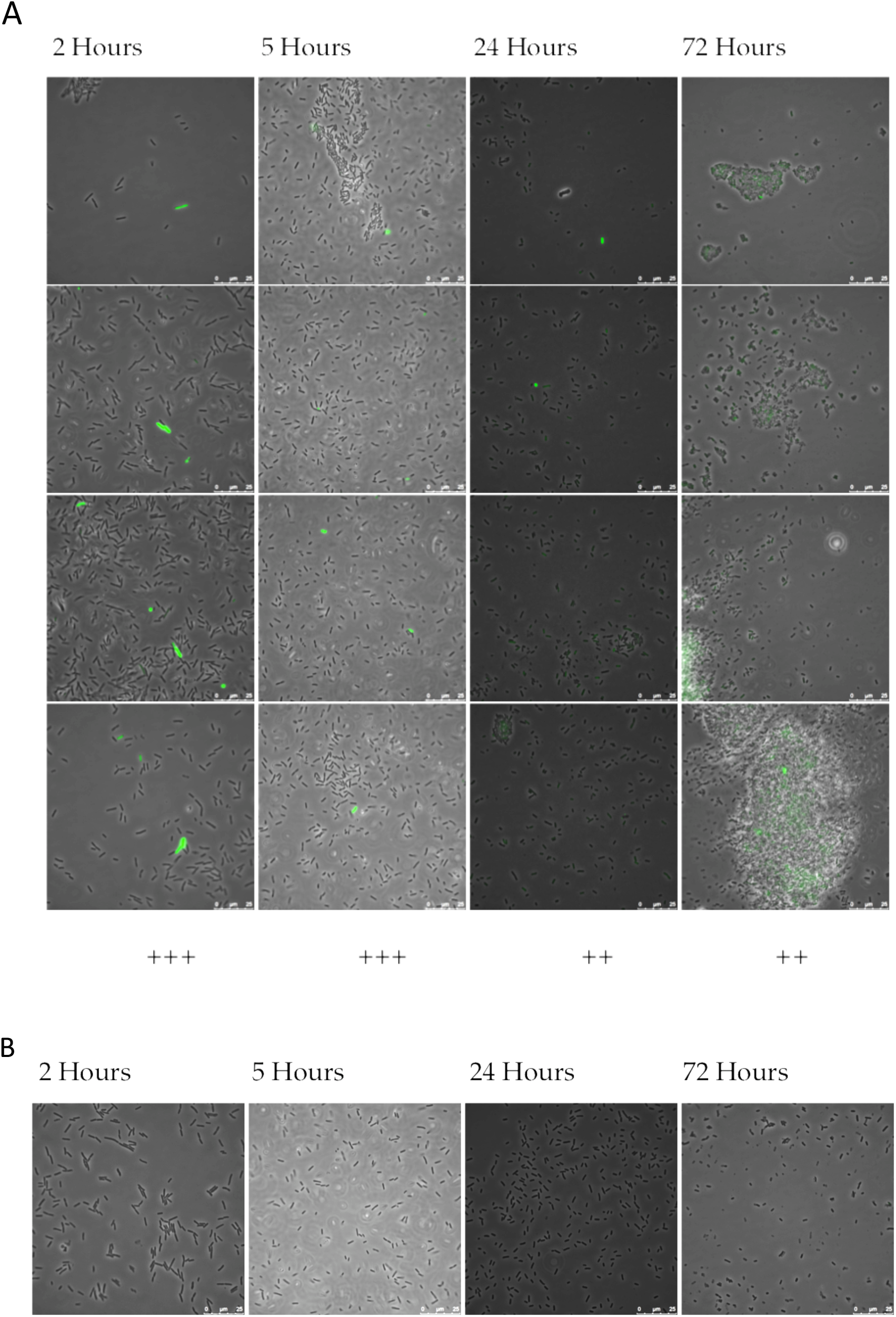
A representative selection of images for 4 time points, for *P. asymbiotica* PB68.1 (*Pa*^PB68^) harbouring **(A)** the *Pa*^PB68^ PVC*pnf pvc1* promoter fusion construct or **(B)** pAGAG negative control reporter plasmid with no promoter. For **(A)** quadruplicate images are displayed vertically as representative of the whole slide sample. Key to qualitative fluorescence indication: “-” is no fluorescence, “++” is low level fluorescence in many cells or a few brighter cells, “+++” is intermediate to high fluorescence in almost all cells, or some very bright isolated cells.

## References

1. Forst, S., et al., Xenorhabdus and Photorhabdus spp.: bugs that kill bugs. Annu Rev Microbiol, 1997. 51: p. 47–72.

2. Ciche, T.A., et al., Cell Invasion and Matricide during Photorhabdus luminescens Transmission by Heterorhabditis bacteriophora Nematodes. Appl Environ Microbiol, 2008. 74(8): p. 2275–87.

3. Somvanshi, V.S., et al., A single promoter inversion switches Photorhabdus between pathogenic and mutualistic states. Science, 2012. 337(6090): p. 88–93.

4. Machado, R.A.R., et al., Whole-genome-based revisit of Photorhabdus phylogeny: proposal for the elevation of most Photorhabdus subspecies to the species level and description of one novel species Photorhabdus bodei sp. nov., and one novel subspecies Photorhabdus laumondii subsp. clarkei subsp. nov. Int J Syst Evol Microbiol, 2018. 68(8): p. 2664–2681.

5. Gerrard, J., et al., Human infection with Photorhabdus asymbiotica: an emerging bacterial pathogen. Microbes Infect, 2004. 6(2): p. 229–37.

6. Gerrard, J.G., et al., Nematode symbiont for Photorhabdus asymbiotica. Emerging Infectious Diseases, 2006. 12(10): p. 1562–1564.

7. Gerrard, J.G., et al., Photorhabdus species: bioluminescent bacteria as emerging human pathogens? Emerg Infect Dis, 2003. 9(2): p. 251–4.

8. Gerrard, J.G., R. Vohra, and G.R. Nimmo, Identification of Photorhabdus asymbiotica in cases of human infection. Commun Dis Intell, 2003. 27(4): p. 540–1.

9. Waterfield, N.R., T. Ciche, and D. Clarke, Photorhabdus and a host of hosts. Annu Rev Microbiol, 2009. 63: p. 557–74.

10. ffrench-Constant, R.H., et al., Photorhabdus: Genomics of a pathogen and symbiont. Bacterial Pathogenomics-Print, ed. M.J. Pallen and K.E. Nelson. 2007. 419–439.

11. Waterfield, N.R., P.J. Daborn, and R.H. ffrench-Constant, Genomic islands in Photorhabdus. Trends Microbiol, 2002. 10(12): p. 541–5.

12. Duchaud, E., et al., The genome sequence of the entomopathogenic bacterium Photorhabdus luminescens. Nat Biotechnol, 2003. 21(11): p. 1307–13.

13. Waterfield, N.R., et al., Rapid Virulence Annotation (RVA): Identification of virulence factors using a bacterial genome library and multiple invertebrate hosts (vol 105, pg 15967, 2008). Proceedings of the National Academy of Sciences of the United States of America, 2009. 106(6): p. 2083–2083.

14. Wilkinson, P., et al., Comparative genomics of the emerging human pathogen Photorhabdus asymbiotica with the insect pathogen Photorhabdus luminescens. BMC Genomics, 2009. 10: p. 302.

15. Cai, X., et al., Entomopathogenic bacteria use multiple mechanisms for bioactive peptide library design. (1755–4349 (Electronic)).

16. Bozhuyuk, K.A.J., et al., Natural Products from Photorhabdus and Other Entomopathogenic Bacteria. (0070–217X (Print)).

17. ffrench-Constant, R.H. and D.J. Bowen, Novel insecticidal toxins from nematode-symbiotic bacteria. Cell Mol Life Sci, 2000. 57(5): p. 828–33.

18. Waterfield, N., et al., Oral toxicity of Photorhabdus luminescens W14 toxin complexes in Escherichia coli. Appl. Environ. Microbiol., 2001. 67(11): p. 5017–24.

19. Waterfield, N.R., et al., The tc genes of Photorhabdus: a growing family. Trends Microbiol, 2001. 9(4): p. 185–91.

20. Meusch, D., et al., Mechanism of Tc toxin action revealed in molecular detail. Nature, 2014. 508(7494): p. 61–5.

21. David L. Erickson, N.R.W., Viveka Vadyvaloo, Daniel Long, Elizabeth R. Fischer, Richard ffrench-Constant, and B. Joseph Hinnebusch, Acute oral toxicity of Yersinia pseudotuberculosis to Xenopsylla cheopis: implications for the evolution of flea-borne transmission of Yersinia pestis. in preparation, 2006.

22. Erickson, D.L., et al., Acute oral toxicity of Yersinia pseudotuberculosis to fleas: implications for the evolution of vector-borne transmission of plague. Cell Microbiol, 2007. 9(11): p. 2658–66.

23. Waterfield, N., et al., The insect toxin complex of Yersinia. Adv Exp Med Biol, 2007. 603: p. 247–57.

24. Hares, M.C., et al., The Yersinia pseudotuberculosis and Yersinia pestis toxin complex is active against cultured mammalian cells. Microbiology-Sgm, 2008. 154: p. 3503–3517.

25. Waterfield, N., et al., The Photorhabdus Pir toxins are similar to a developmentally regulated insect protein but show no juvenile hormone esterase activity. FEMS Microbiol Lett, 2005. 245(1): p. 47–52.

26. Ahantarig, A., et al., PirAB Toxin from Photorhabdus asymbiotica as a Larvicide against Dengue Vectors. Applied and Environmental Microbiology, 2009. 75(13): p. 4627–4629.

27. Sirikharin, R., et al., Characterization and PCR Detection Of Binary, Pir-Like Toxins from Vibrio parahaemolyticus Isolates that Cause Acute Hepatopancreatic Necrosis Disease (AHPND) in Shrimp. (1932–6203 (Electronic)).

28. Daborn, P.J., et al., A single Photorhabdus gene, makes caterpillars floppy (mcf), allows Escherichia coli to persist within and kill insects. Proceedings of the National Academy of Sciences of the United States of America, 2002. 99(16): p. 10742–10747.

29. Waterfield, N.R., et al., The insecticidal toxin makes caterpillars floppy 2 (Mcf2) shows similarity to HrmA, an avirulence protein from a plant pathogen. Fems Microbiology Letters, 2003. 229(2): p. 265–270.

30. Dowling, A.J., et al., The Mcf1 toxin induces apoptosis via the mitochondrial pathway and apoptosis is attenuated by mutation of the BH3-like domain. Cellular Microbiology, 2007. 9(10): p. 2470–2484.

31. Yang, G., et al., Photorhabdus virulence cassettes confer injectable insecticidal activity against the wax moth. J Bacteriol, 2006. 188(6): p. 2254–61.

32. Ghequire, M.G. and R. De Mot, The Tailocin Tale: Peeling off Phage Tails. Trends Microbiol, 2015. 23(10): p. 587–90.

33. Hapeshi, A. and N.R. Waterfield, Photorhabdus asymbiotica as an Insect and Human Pathogen. Curr Top Microbiol Immunol, 2017. 402: p. 159–177.

34. Hurst, M.R., T.R. Glare, and T.A. Jackson, Cloning Serratia entomophila antifeeding genes--a putative defective prophage active against the grass grub Costelytra zealandica. J Bacteriol, 2004. 186(15): p. 5116–28.

35. Heymann, J.B., et al., Three-dimensional structure of the toxin-delivery particle antifeeding prophage of Serratia entomophila. J Biol Chem, 2013. 288(35): p. 25276–84.

36. Durand, E., et al., Biogenesis and structure of a type VI secretion membrane core complex. Nature, 2015. 523(7562): p. 555–60.

37. Shikuma, N.J., et al., Marine tubeworm metamorphosis induced by arrays of bacterial phage tail-like structures. Science, 2014. 343(6170): p. 529–33.

38. Sarris, P.F., et al., A phage tail-derived element with wide distribution among both prokaryotic domains: a comparative genomic and phylogenetic study. (1759–6653 (Electronic)).

39. Wilkinson, P., et al., New plasmids and putative virulence factors from the draft genome of an Australian clinical isolate of Photorhabdus asymbiotica. FEMS Microbiol Lett, 2010. 309(2): p. 136–43.

40. Belogurov, G.A., et al., Functional specialization of transcription elongation factors. EMBO J, 2009. 28(2): p. 112–22.

41. Mulley, G., et al., From Insect to Man: Photorhabdus Sheds Light on the Emergence of Human Pathogenicity. PLoS One, 2015. 10(12): p. e0144937.

42. Vernikos, G.S. and J. Parkhill, Interpolated variable order motifs for identification of horizontally acquired DNA: revisiting the Salmonella pathogenicity islands. Bioinformatics, 2006. 22(18): p. 2196–203.

43. Leiman, P.G., et al., Morphogenesis of the T4 tail and tail fibers. Virol J, 2010. 7: p. 355.

44. Russell, A.B., S.B. Peterson, and J.D. Mougous, Type VI secretion system effectors: poisons with a purpose. Nat Rev Microbiol, 2014. 12(2): p. 137–48.

45. Knust, Z. and G. Schmidt, Cytotoxic Necrotizing Factors (CNFs)-A Growing Toxin Family. Toxins (Basel), 2010. 2(1): p. 116–27.

46. Kapitein, N. and A. Mogk, Deadly syringes: type VI secretion system activities in pathogenicity and interbacterial competition. Curr Opin Microbiol, 2013. 16(1): p. 52–8.

47. Taylor, N.M.I., M.J. van Raaij, and P.G. Leiman, Contractile injection systems of bacteriophages and related systems. Mol Microbiol, 2018. 108(1): p. 6–15.

48. Hurst, M.R., et al., Isolation and characterization of the Serratia entomophila antifeeding prophage. FEMS Microbiol Lett, 2007. 270(1): p. 42–8.

49. Hurst, M.R., M. O’Callaghan, and T.R. Glare, Peripheral sequences of the Serratia entomophila pADAP virulence-associated region. Plasmid, 2003. 50(3): p. 213–29.

50. Carter, H.D., V. Svetlov, and I. Artsimovitch, Highly divergent RfaH orthologs from pathogenic proteobacteria can substitute for Escherichia coli RfaH both in vivo and in vitro. J Bacteriol, 2004. 186(9): p. 2829–40.

51. Tobias, N.J., et al., Photorhabdus-nematode symbiosis is dependent on hfq-mediated regulation of secondary metabolites. (1462–2920 (Electronic)).

52. Joyce, S.A. and D.J. Clarke, A hexA homologue from Photorhabdus regulates pathogenicity, symbiosis and phenotypic variation. Mol Microbiol, 2003. 47(5): p. 1445– 57.

53. Meijer, M., et al., Nucleotide sequence of the origin of replication of the Escherichia coli K-12 chromosome. Proc Natl Acad Sci U S A, 1979. 76(2): p. 580–4.

54. Ge, P., et al., Atomic structures of a bactericidal contractile nanotube in its pre- and postcontraction states. Nat Struct Mol Biol, 2015. 22(5): p. 377–82.

55. Reynolds, S.E. and S.F. Nottingham, Food and water economy and its relation to growth in the 5th-instar larvae of the Tobacco Hornworm, Manduca sexta. Journal of Insect Physiology, 1985. 31: p. 119–127.

56. Thanwisai, A., et al., Diversity of Xenorhabdus and Photorhabdus spp. and Their Symbiotic Entomopathogenic Nematodes from Thailand. PLoS ONE, 2012. 7(9).

57. Jacobi, C.A., et al., In vitro and in vivo expression studies of yopE from Yersinia enterocolitica using the gfp reporter gene. Mol Microbiol, 1998. 30(4): p. 865–82.

58. Kanter-Smoler, G., A. Dahlkvist, and P. Sunnerhagen, Improved method for rapid transformation of intact Schizosaccharomyces pombe cells. Biotechniques, 1994. 16(5): p. 798–800.

59. Brillard, J., et al., The PhlA hemolysin from the entomopathogenic bacterium Photorhabdus luminescens belongs to the two-partner secretion family of hemolysins. J Bacteriol, 2002. 184(14): p. 3871–8.

60. Schmidt, G., et al., Gln 63 of Rho is deamidated by Escherichia coli cytotoxic necrotizing factor-1. Nature, 1997. 387(6634): p. 725–9.

61. Sugai, M., et al., Cytotoxic necrotizing factor type 2 produced by pathogenic Escherichia coli deamidates a gln residue in the conserved G-3 domain of the rho family and preferentially inhibits the GTPase activity of RhoA and rac1. Infect Immun, 1999. 67(12): p. 6550–7.

62. Schmidt, G., et al., Identification of the C-terminal part of Bordetella dermonecrotic toxin as a transglutaminase for rho GTPases. J Biol Chem, 1999. 274(45): p. 31875–81.

63. Muggeo, V.M. and G. Adelfio, Efficient change point detection for genomic sequences of continuous measurements. Bioinformatics, 2011. 27(2): p. 161–6.

64. Sievers, F., et al., Fast, scalable generation of high-quality protein multiple sequence alignments using Clustal Omega. Mol Syst Biol, 2011. 7: p. 539.

